# Statistical modelling of bacterial promoter sequences for regulatory motif discovery with the help of transcriptome data: application to *Listeria monocytogenes*

**DOI:** 10.1101/723346

**Authors:** Ibrahim Sultan, Vincent Fromion, Sophie Schbath, Pierre Nicolas

## Abstract

Automatic *de novo* identification of the main regulons of a bacterium from genome and transcriptome data remains a challenge. To address this task, we propose a statistical model of promoter DNA sequences that can use information on exact positions of the transcription start sites and condition-dependent expression profiles. Two main novelties are to allow overlaps between motif occurrences and to incorporate covariates summarising expression profiles (e.g. coordinates in projection spaces or hierarchical clustering trees). All parameters are estimated using a dedicated trans-dimensional Markov chain Monte Carlo algorithm that adjusts, simultaneously, for many motifs and many expression covariates: the width and palindromic properties of the corresponding position-weight matrices, the number of parameters to describe position with respect to the transcription start site, and the choice of relevant expression covariates. A data-set of transcription start sites and expression profiles available for the *Listeria monocytogenes* is analysed. The results validate the approach and provide a new global view of the transcription regulatory network of this important model food-borne pathogen. A previously unreported motif that may play an important role in the regulation of growth was found in promoter regions of ribosomal protein genes.

## Introduction

Motif discovery in DNA sequences is an old problem of bioinformatics for which many approaches have been developed [1, 2] but automatic *de novo* identification of the main regulons of an organism as “simple” as a bacterium remains a challenge. The representation chosen for the motifs is at the heart of the methodology. Word-based methods usually represent motifs as consensus strings written in an alphabet allowing degenerate symbols and rely, for scoring, on an hypothesis testing framework to detect deviation from a null hypothesis, such as Markov sequence [3] or equal occurrence frequencies between coexpression clusters [4, 5]. Position-Weight Matrices (PWMs) are more precise description of motifs that can account for the expected frequencies of the 4 DNA nucleotides (`A,C,G,T`) at each position within the motif. Beside this probabilistic representation of the motif, a full probabilistic model involves also a model for the background sequence (outside motif occurrences) and a model for the positions of motif occurrences in the sequence set. Motif discovery is then cast as the problem of estimating PWMs. The first algorithms implementing these ideas [6, 7] remain among the most powerful and widely used tools to search for motifs based only on the nucleotide composition properties. Due to motif degeneracy and limited number of occurrences, these approaches are usually successful only when sequence data-sets enriched for particular motifs can be defined, most often based on experimental data. Hence, transcriptional regulatory network reconstruction tends to be an incremental process in which new components of the network are added one by one.

Chromatin immunoprecipitation (ChIP) is the experimental technique that had the deepest impact on the field of motif discovery [2]. Nevertheless, the *a priori* selection of combinations of transcription factors (TFs) and biological conditions is an intrinsic limitation of ChIP for *de novo* motif discovery at a system level. Furthermore, a ChIP experiment requires either a specific antibody to recognise the TF or genetic engineering of a functionally active tagged TF. In parallel to ChIP, microarrays and RNA-Seq have been extensively applied to compare gene expression between growth conditions and/or specific mutants and results have been early and widely used for bacterial transcriptional network reconstruction [8, 9]. With the goal to unravel globally the transcription regulatory network of a bacterium, expression profiles across many conditions can be collected without focusing on specific TFs and without genetic engineering [10, 11]. Genome-wide maps of transcription start sites (TSSs) constitute another type of transcriptome data which allows to focus the search for regulatory motifs and is increasingly available for bacteria [12, 13].

The development of specific methods for *de novo* motif discovery using expression profiles has attracted less attention than the use of ChIP data. The task is also more difficult because the data are not directly connected to a specific TF–DNA interaction. Nevertheless, a diversity of approaches has been proposed based on different methodological concepts such as: mutual information in FIRE [14], enrichment test in GEMS [4], or regression of expression data *y* given sequence data *x* in REDUCE [15] and MatrixREDUCE [16]. The first two approaches involve transforming the expression data into one-dimensional categorical values, while the third approach can directly accommodate multidimensional continuous expression data. The algorithm implemented in RED2 [5] intends to bypass the need for clustering of the first two approaches by applying enrichment tests or computing mutual information for overlapping sets of genes that are close in the expression space (neighbourhoods). These approaches face the difficulty to find appropriate summary or probabilistic models for expression data.

The alternative viewpoint adopted in this work consists in modelling the sequence data *x* given the expression data *y*. A key benefit of this choice is to build directly upon the powerful sequence modelling approaches for *de novo* motif discovery based on PWMs and full probabilistic modelling of the sequences, establishing a continuum between discovery of motifs related and unrelated to the available expression data. This makes it possible to envision the simultaneous use of the expression data and of all the statistical properties of the sequence. In [10], we already proposed an approach for the discovery of Sigma factor binding sites based on modelling *x*|*y*. The generality of the model was however limited since it was specifically tailored for sigma factor binding sites whose specificity is to delineate and partition the promoter space [17]. Regulation by sigma factors tends thus to correspond to a preponderant and non-overlapping level of regulation that is particularly well captured by the structure of a hierarchical clustering tree and did not justify to model the occurrence of more than one motif per sequence [10]. Incorporation of positional information proved helpful in the case of sigma factor binding sites [10] whose positions are strongly constrained with respect to the TSS.

This work develops a coherent probabilistic model of the DNA sequence to address the task of automatic *de novo* identification of the main regulons (not restricted to sigma factors) of a bacterium from genome and transcriptome data. Two main novelties of the proposed model are to allow overlaps between motif occurrences and to incorporate covariates summarising expression profiles into the probability of occurrence in a given promoter region. Covariates can correspond to the positions of the genes on an axis such as obtained by PCA [18] or ICA [19, 20] but we also show how to use positions in a hierarchical clustering trees [21, 8]. All the parameters are estimated in a Bayesian framework using a dedicated trans-dimensional MCMC algorithm. In order to validate the approach, we applied it to the food-borne pathogen *L. monocytogenes* on which a wealth of transcriptomics data has been collected owing to its status as important human pathogen and model bacterium for the study of host-pathogen interaction and bacterial transcriptomics [22]. Sources of transcriptome data include a landmark study using RNA-Seq and high-density tiling arrays [23], an early use of genome-wide TSS mapping [24], and a comprehensive work done to aggregate available transcriptome data-sets in a single database [25].

## Methods and data

### Probabilistic model of promoter sequences

#### Model overview

The sequence data considered here consists of 𝒩 DNA sequences of same length ℒ. Let denote by *x* = (*x*_*n*_)_*n*=1:𝒩_ this set of sequences and *x*_*n,l*_ ∈ {`A, C, G, T`} the nucleotide in position *l* of the *n*^th^ sequence. Probabilistic representation of these sequences involves ℳ + 1 unobserved components that capture the heterogeneity of nucleotide composition. These components consist of ℳ motif models (PWMs), denoted by (*θ*_1_, …, *θ* _ℳ_), with respective widths (*w*_1_, …, *w* _ℳ_) and a background Markov model of order *v* whose parameters are denoted by *θ*_0_. All the ℳ motifs are simultaneously searched for. For simplicity, the model assumes <u>z</u>ero or <u>o</u>ne <u>o</u>ccurrence per sequence (ZOOPS, [6]) of each motif but occurrences of different motifs are allowed and may even overlap. The occurrences of the different motifs are modelled as mutually independent.

Statistical inference is carried out in a Bayesian framework and the parameters are thus treated as random variables drawn from prior distributions. The Directed Acyclic Graph (DAG) of the model is shown in Fig 1 and its complete mathematical presentation is found in S1 Appendix. We explain below the purpose of the different variables with a focus on the most salient characteristics of the model.

**Figure 1:**
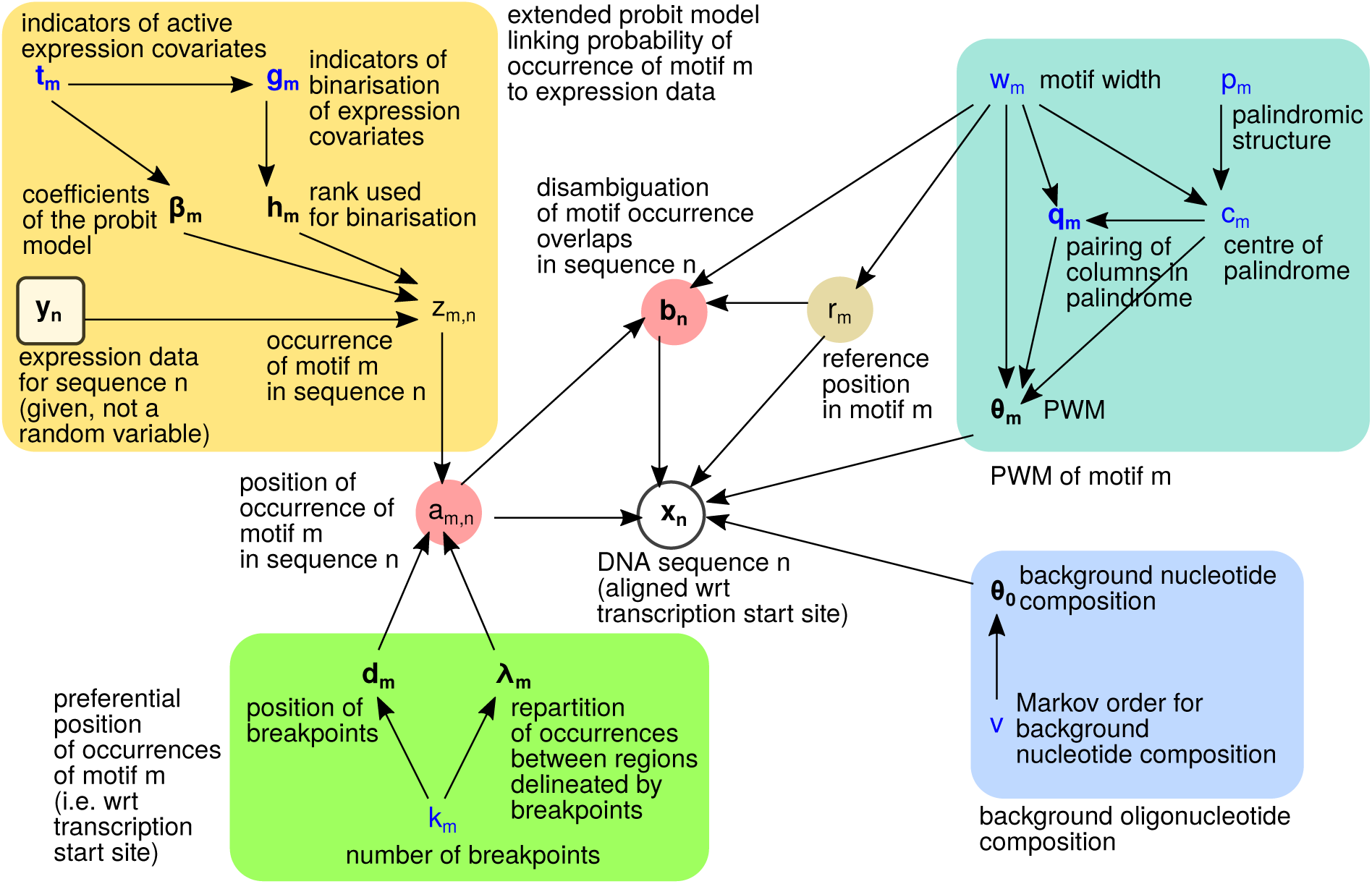
Hierarchical model of promoter sequences. DAG where vertices represent the variables (parameters, hidden variables, observed data) and edges show the factorisation of the joint probability distribution. Coloured areas highlight groups of variables that contribute to a same ingredient of the model (e.g. yellow-ochre for incorporation of expression data). In this representation we show the variables for only one motif *m* and one sequence *n*. Multidimensional variables (vectors or matrices) are boldfaced; variables whose values change the dimension of the model are in blue; continuous random variables are denoted by Greek letters.

The observed sequence data *x* is at the centre of this DAG. Connected to *x, a* = (*a*_*m,n*_)_*m*=1: ℳ, *n*=1:𝒩_ is a layer of random variables that encode the position of the occurrences of the motifs, with *a*_*m,n*_ ∈ {0, 1, …, ℒ} being the position of motif *m* in sequence *n*; 0 encodes the absence of occurrence. For practical reasons pertaining to the implementation of the update of the motif width *w*_*m*_ in the MCMC algorithm we do not systematically rely on the first position within motif *m* to record its position. Instead, we use the reference position *r*_*m*_ ∈ {1, …, *w*_*m*_} which allows motif occurrences to extend somewhat outside of the sequences.

#### Incorporating expression data as covariates

Information from expression data is incorporated into the probability of occurrence of a motif via a probit regression framework. Implementation of this model relies on a data augmentation scheme [26] that introduces a Gaussian latent variable (*z*_*m,n*_).

Let’s consider the availability of a number *𝒞* of vectors of continuous variables denoted by *y* = (*y*_*n,c*_)_*n*=1: 𝒩, *c*=1: *𝒞*_ that can be used as covariates. The standard probit model would write the probability of occurrence of motif *m* in sequence *n* as

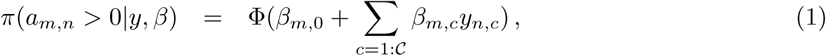

where Φ is the cumulative distribution function of the standard normal distribution and *β* contains the regression coefficients (*β*_*m,c*_ ∈ ℝ).

We developed an extension for this model that allows binarisation of expression covariates according to automatically adjusted cut-offs. This extension is very appealing here because it can account for sharp switches in the probability of occurrence as a function of the position of the sequence in the expression space; exactly like if co-expression clusters defined on the basis of gene coordinates on axis *c* were also entered as covariates. Importantly, these sharp switches do not imply a probability of occurrence jumping between 0 and 1 as when increasing |*β*_*m,c*_|in the standard probit.

To allow automatic selection of the covariates relevant for the occurrences of motif *m* and their binarisation (both choices impacting on model dimension), the model involves the following variables,

- *t*_*m*_ = (*t*_*m*,1_, …, *t*_*m, 𝒞*_) where *t*_*m,c*_ ∈ {0, 1} indicates whether covariate *c* should be taken into account;
- *β*_*m*_ = (*β*_*m*,0_, *β*_*m*,1_, …, *β*_*m, 𝒞*_) where *β*_*m, c*_ ∈ ℝ represents, when *t*_*m,c*_ = 1, a coefficient specific to covariate *c, β*_*m*,0_ being the intercept parameter;
- *g*_*m*_ = (*g*_*m*,1_, …, *g*_*m, 𝒞*_) where *g*_*m, c*_ ∈ {0, 1} indicates, when *t*_*m,c*_ = 1, whether the values in the vector *y*_·,*c*_ are binarised;
- *h*_*m*_ = (*h*_*m*,1_, …, *h*_*m, 𝒞*_) where *h*_*m,c*_ ∈ {1, …, 𝒩 − 1} indicates, when *t*_*m,c*_ = 1 and *g*_*m,c*_ = 1, the rank in the vector *y*_·,*c*_ of the value used for binarisation; 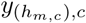 denotes the corresponding cut-off.

In keeping with the probit regression framework, the probability of occurrence of motif *m* in sequence *n* writes then as

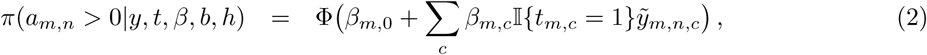

where 𝕀 is the indicator function (𝕀{*z*} = 1 if *z* is true, 0 otherwise) and 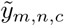 corresponds to *y*_*n,c*_ after taking into account a possible binarisation (see S1 Appendix for how binarised values depend on *h*_*m,c*_). This model can capture a great diversity of relationships between expression covariates and probability of motif occurrence (Fig 2ABC).

**Figure 2:**
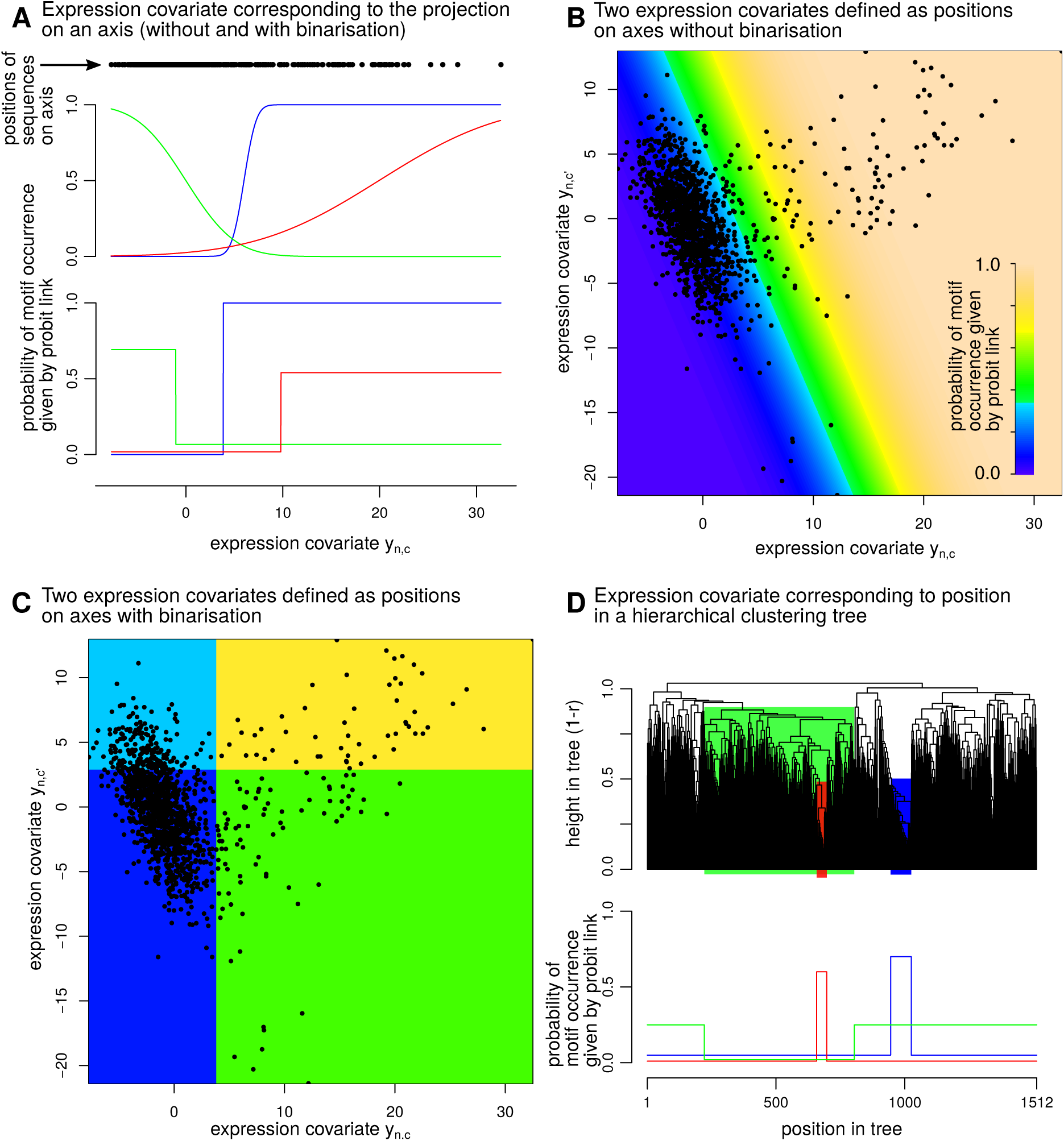
Graphical illustration of the use of the extended probit model. A. Linking probability of motif occurrence to one expression covariate. Upper plot: classical probit model. Lower plot: probit model with covariate binarisation. Three sets of regression coefficients are illustrated for each case. **B** Linking probability of motif occurrence to two expression covariates (one set of regression coefficients). **C** Linking the probability of motif occurrence to two expression covariates with binarisation (one set of regression coefficients). **D** Linking probability of motif occurrence to position in a tree (three sets of regression coefficients).

#### Handling trees as covariates

Further extending the model described by Eq (2), we consider that covariates can come not only as vectors of continuous values but also in the form of trees. Indeed, the binarisation proposed above makes it also possible to incorporate whole tree structures in the regression model. In this case, binarisation involves the choice of node instead of a cut-off value along an axis.

Here, trees are obtained from expression data by usual techniques of hierarchical clustering. They are thus rooted, binary, with all leaves at a same distance from the root. Such a tree covariate *c*, can be conveniently encoded (topology and branch lengths) as a vector of 𝒩 − 1 internal node heights and a *s*_*c*_(𝒩 − 1) × 2 matrix that identifies the pair of subtrees merged at each internal node that, together, replace the vector *y*_*c*_ when the covariate *c* is a tree. Choice of a node allows to associate different probabilities of motif occurrence inside and outside the hanging sub-tree (Figure 2D).

#### Modelling occurrence positions

DNA sequences are aligned with respect to experimentally determined TSSs such that the position of motif occurrence encoded in *a*_*m,n*_ corresponds to a precise position relative to the TSS.

Given an occurrence of motif *m* in sequence *n* (event {*a*_*m,n*_ *>* 0} modelled by the extended probit), the probability density function of its exact position is a piece-wise constant function defined by *k*_*m*_ breakpoints located at 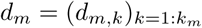 and the probabilities 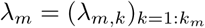 to find the occurrence between each of these breakpoints.

#### Allowing motif occurrences to overlap

The model allows motif occurrences to overlap. This feature underpins the simplifying assumption of mutual independence between the occurrences of the ℳ motifs made when modelling the position of the motif with respect to the TSS and the link with expression data. Another key benefit of allowing overlap between motif occurrences is to allow the update motif width (*w*_*m*_) without having to avoid collisions of motif occurrences.

When *o*_*n,l*_ motifs overlap position *l* in sequence *n, x*_*n,l*_ is modelled as drawn from the arithmetic mean of PWM columns, namely

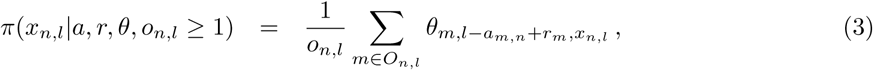

where *O*_*n,l*_ denotes the set of motifs that overlap position *l*, and *θ*_*m,w,u*_ is the probability of nucleotide *u* at position *w* of motif *m*. The resulting density can be seen as the marginal of an equal-weight mixture. In keeping with the usual data augmentation scheme for Bayesian inference of mixture models [27], a layer of latent random variables (denoted by *b*) is introduced to encode the disambiguation of the overlaps at each position (*n, l*) of the sequence set.

#### Palindromic motifs

Dimeric nature and symmetry of many TFs explain that many of the known binding motifs are palindromic in the sense that symmetric positions with respect to the centre of the motif appear as mirrored according to Watson-Crick base pairing rule (`A : T` and `C : G`). Palindromic constraints on PWMs reduce the dimension of the model, increasing the amount of data available to estimate each parameter and decreasing the size of the search space for the parameter values.

Instead of imposing a strict palindromic constraint on all or a subset of the motifs, we developed a more flexible representation that allows, for each motif *m*, intermediate states between non-palindromic and fully-palindromic structures. The following set of variables define the active constraints on 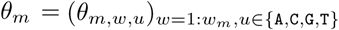:

- *p*_*m*_, a binary variable indicating if the motif *m* has a (possibly partial) palindromic structure,
- *c*_*m*_ ∈ {1.5, 2, 2.5, …, *w*_*m*_ − 0.5}, a discrete variable recording the position of the centre of symmetry of the palindromic structure; the range for *c*_*m*_ allows two types of palindromic structure (“even” and “odd”, where the odd type contains a central unpaired column at *w* = *c*_*m*_);
- *q*_*m,w*_, a binary variable to indicate whether columns *w* and 2*c*_*m*_ − *w* are paired, i.e. 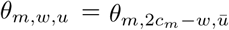 when 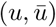 is a Watson-Crick pair.

There are several motivations for this representation. First, identifying as many motifs as possible simultaneously is incompatible with having all the motifs palindromic. Second, intermediate states allow to design algorithms that gradually increase or decrease the number of free parameters. Finally, while some symmetry is often obvious, it may be unclear to which extent a biological motif is fully palindromic or partially palindromic.

#### Bayesian inference

The priors used intend to be non-informative and a Markov Chain Monte Carlo (MCMC) algorithm was built to sample the joint posterior

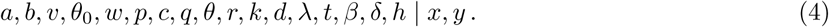

To cope with the high dimension of the target, the algorithm is a block MCMC sampler [28] composed of 14 types of steps designed to update separate subsets (blocks) of variables. A sweep combines the different steps. Dimension changes rely on the Reversible-Jump methodology [29]. They are needed to update the Markov order of the background (variable *v*), the active covariates and their possible binarisation (block *t*_*m,c*_, *g*_*m,c*_, *h*_*m,c*_), and the variables encoding the palindromic structure of the motif (block *p*_*m*_, *c*_*m*_, *q*_*m*_). Under circumstances where the probability distribution of the variables whose dimension is modified can be integrated-out, the Reversible-Jump can be done in a Gibbs manner, i.e. by direct sampling from the conditional distribution. The algorithm proceeds this way for the joint update of (*t*_*m,c*_, *g*_*m,c*_, *h*_*m,c*_) and the joint update of (*p*_*m*_, *c*_*m*_, *q*_*m*_). Correctness of the MCMC algorithm was carefully checked by implementing a successive-conditional simulator to reveal analytical and coding errors [30]. The MCMC algorithm is described in S1 Appendix and has been implemented in C++ in the program Multiple.

### Data set for application to *Listeria monocytogenes*

#### Transcription start sites and expression profiles

To define promoter sequences and to align them with respect to TSSs, we used the repertoire of 2,299 TSSs mapped at 1 bp resolution on *L. monocytogenes* EGDe genome sequence [24]. Promoter sequences were defined as the 121 bp spanning from position -100 to +20 relative to each TSS, in keeping with the size of the regions that we previously found to be enriched for the presence of known TF binding sites [11]. To remove promoter sequence overlaps on a same strand, we used a simple greedy procedure that incorporated non-overlapping promoters one-by-one in the order of decreasing read-count (reflecting level of experimental support and transcriptional activity for the TSS [24]). This led to a set of 1,545 non-overlapping promoter regions (67% of the initial list of TSSs).

For the expression data, we relied on the compendium data-set established by aggregation of many different studies to build the listeriomics website [25]. As downloaded, the data had dimensions 3,159 ×254, where each row corresponds to a gene of *L.monocytogenes* EGDe and each column corresponds to the log of an expression ratio (log fold-change) comparing two samples (mutants, growth conditions, strains …) from a same study. Some columns and rows contained many missing values due to the heterogeneity of technologies. The number of columns was reduced from 254 to 165 after discarding the columns with a number of missing values higher than 1.5 times the median. In parallel, the number of rows was reduced from 3,159 to 2,825 based on the same criterion. Finally, the gene name associated with each TSS [24] permitted to match 1,512 out of 1,545 non-overlapping promoter sequences to one of these 2,825 genes with expression data. This resulted in an expression matrix of dimension 1,512 ×165 whose 165 columns represented 31 published expression studies, and included 17 RNA-Seq and 25 tiling array profiles.

#### Building covariates for motif discovery

Both hierarchical clustering and projection methods were used as dimension reduction techniques to summarise the expression matrix. Two types of hierarchical clustering were applied corresponding to different options of `hclust` function provided by `R stats` package: Ward and average-link based on Pearson distance (1 −*r*, where *r* is Pearson correlation coefficient). The distances between genes needed to build the trees were computed after centring and scaling of the rows (genes) of a symmetric expression matrix obtained by duplicating each column with a negative sign to remove the effect of the arbitrary orientation of the log fold-changes.

As projection methods, we applied PCA and ICA implemented in functions `prcomp` (package `stats`) and `fastICA` [31] of `R` without scaling the expression data. For PCA, we kept the 20 first components of the PCA (accounting for 68.8% of the total variance) after examining the rate of decrease of the residual variance. When applying ICA, the target dimension (number of components) is fixed before numerical optimisation and the algorithm can converge to different projections that share only a subset of components. In keeping with the idea developed by [20], we choose the target dimension *K* = 40 after examining the stability of the components and only stable components were used. Here, we used average-link clustering based on absolute Pearson correlation coefficient (*r*) between columns of the source matrix (dimension 1,512 ×K) and a cut-off |*r*| ≥ 0.8 to compare components between runs. The algorithm was run 100 times which lead to 26 components found in at least 80% of the runs.

The final set of 50 covariates used in the motif discovery analysis consisted of the 20 first PCA components (covariates numbered 1 to 20), the 26 stable ICA components (covariates numbered 21 to 46), and the hierarchical clustering trees obtained by Ward and average-link methods (covariates numbered 47 and 48). The two trees were further duplicated (covariates 49 and 50) to make it possible for the model to use two nodes of the same tree.

## Results

### Exploration of the posterior landscape

Dynamics of evolution of the parameters describing the ℳ motif components during MCMC runs confirmed that the algorithm was able to adjust simultaneously the characteristics of many PWMs. To illustrate the algorithm behaviour, Fig 3A depicts the parallel evolution in a same MCMC run of two PWMs in terms of width, nucleotide composition, and palindromic structure.

**Figure 3:**
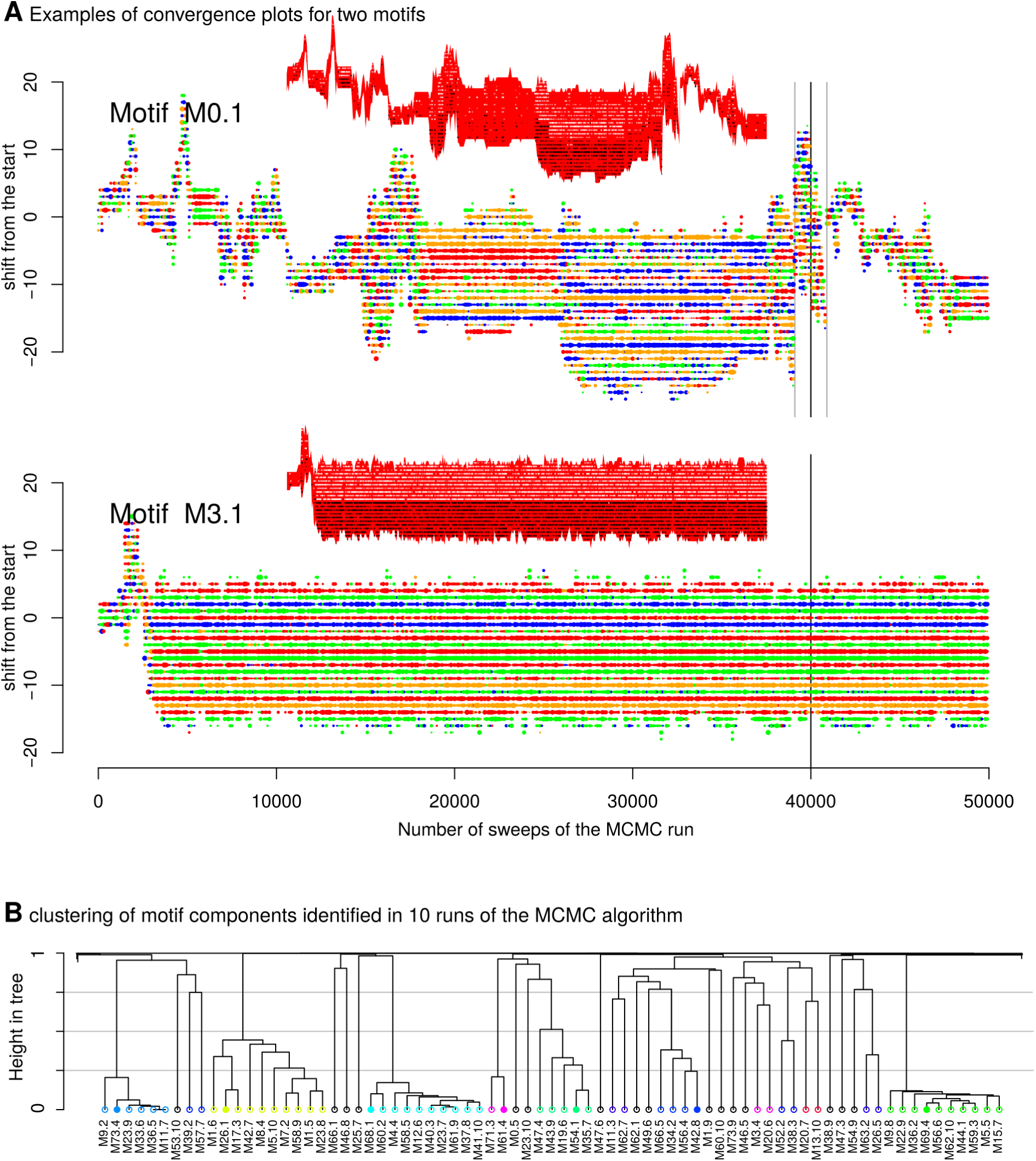
Exploration of the posterior landscape. **A** Convergence plots for two motifs across the 50,000 sweeps of the algorithm. Coloured dot points correspond to PWM columns, colour indicates the most likely nucleotide (green for `A`, blue for `C`, orange for `G`, red for `T`) and diameter reflects the information content of the nucleotide frequency. Increase or decrease of the width of the PWM on the 5’- and 3’-sides are represented by change in the number of coloured dots (on lower and upper side of the coloured area, respectively). A black vertical line is drawn at sweep number 40,000 (end of our burn-in period). Grey vertical lines indicate sweeps at which the y-coordinate was recentred for the purpose of the representation. Evolution of the palindromic structure is represented in insert plots (at lower scale) where the evolution of PWM width is represented by a red area and paired columns are indicated by black and white dots. **B** Hierarchical clustering of motif components to extract stable motifs. Only a fraction of the tree is represented here (75 motif components out of 750). High-level clusters (defined by height 0.75) are represented here by different colours. Motifs belonging to the final list of 40 representative stable motifs are indicated by filled circles.

For *de novo* motif discovery, it was important to show that stability of a motif component across tens of thousands of MCMC sweeps as seen in Fig 3A was not caused by slow-mixing but truly reflected attraction to peaks of the posterior density and should therefore be treated as relevant motif predictions. Since reaching similar motif components independently from different starting points is indicative of the second scenario, we performed 10 independent parallel runs of the MCMC algorithm. Each run consisted of 50,000 MCMC sweeps from a random starting point ℳ and was fixed to 75 (≈ 4 weeks of CPU time). Only the last 10,000 sweeps were used in our analysis to characterise the posterior distributions after recording the values of the variables every 100 sweeps.

The 10 runs produced information on 750 (10 × 75) motif components that were named from M1.0 (random seed 1, motif 0) to M10.74. We analysed and compared these motif components to extract distinct well supported motifs that were not only stable across the last 10,000 sweeps but also found in at least two runs. To declare that two motif components correspond in fact to the same motif, we compared the sets of positions in the sequences that were predicted to be covered by the occurrences of each motif which summarise the information of the different ingredients of the motif model (PWM, preferential position with respect to TSS, link with expression data). Namely, we considered the positions with estimated posterior probability of coverage greater than 0.5 and the pairwise distance between two motif components *i* and *j* was computed as

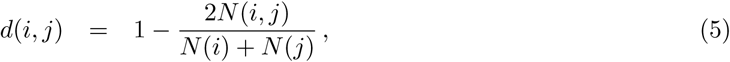

where *N* (*i*) and *N* (*j*) denote the total number of positions covered by each motif and *N* (*i, j*) is the number of positions covered by both motifs. With this formula, *d*(*i, j*) = 0 if the positions covered are exactly the same and *d*(*i, j*) = 1 if they do not overlap. A special case of no overlap corresponds to *N* (*i*) = 0 which concerned 190 of the 750 motifs and typically reflected instability during the last 10,000 sweeps. The minimal distance that we observed between two motif components within the same run was and 92.7% of the motif components where at distance ≥0.9 of any other motif components from the same run. This shows that allowing motif occurrences to overlap in the model is an effective strategy to enforce the convergence of motif components to distinct motifs.

The list of distinct well supported motifs was established after average-link hierarchical clustering of the 750 motifs based on the distance defined in Eq (5). This tree whose a section is shown in Fig 3B was cut at two different heights: 0.75 to define high-level clusters that separate well-distinct motifs; 0.25 to define low-level clusters containing very similar motifs. In each high-level cluster containing at least one low-level cluster, a representative motif was selected as the member of the low-level clusters with the highest *N* (*i*). This procedure defined the set of 40 representative motifs reported in Table 1.

**Table 1:** Summary of the 40 stable motifs identified in *L. monocytogenes* EGD-e.

| motif<sup>a</sup> | #<sup>b</sup> | #(0.8-0.2)<sup>c</sup> | W<sup>d</sup> | IC1<sup>e</sup> | Pal<sup>f</sup> | Pos<sup>g</sup> | FC<sup>h</sup> | Runs<sup>i</sup> | Comment<sup>j</sup> |
| --- | --- | --- | --- | --- | --- | --- | --- | --- | --- |
| M71.2 | 1,325 | 1,174-1,435 | 9 | 3 | 1.3 | -9 [1] | 0.44 | 7:10 | SigA -10 |
| M71.8 | 1,308 | 848-1,506 | 6 | 2 | 1.6 | 1 [1] | 0.46 | 13:16 | TSS (A) |
| M36.4 | 1,033 | 552-1,382 | 6 | 3 | 0 | -32 [2] | 0.48 | 10:10 | SigA -35 (TTG) |
| M55.7 | 836 | 117-1,486 | 5 | 0 | 1.7 | -25 [40] | 0.58 | 3:7 | SigA -10 extension (CT) |
| M8.3 | 792 | 331-1,273 | 5 | 1 | 0 | 4 [1] | 0.53 | 10:15 | TSS (G) |
| M9.10 | 735 | 447-1,165 | 8 | 4 | 0.6 | -13 [1] | 0.50 | 7:8 | SigA (extended -10, TG) |
| M61.5 | 561 | 171-1,153 | 5 | 5 | 0 | 14 [10] | 0.41 | 6:8 | RBS (GGAGG) |
| M59.9 | 297 | 96-844 | 21 | 7 | 0 | -60 [42] | 0.36 | 4:10 | T-rich element |
| M26.1 | 252 | 107-640 | 13 | 5 | 2.1 | -79 [23] | 0.73 | 6:10 | SigA -10 on reverse strand |
| M27.6 | 240 | 153-590 | 6 | 5 | 0.4 | -33 [2] | 0.48 | 6:6 | SigA -35 (TTGAC) |
| M29.8 | 122 | 91-150 | 12 | 4 | 2.2 | -13 [2] | 3.06 | 10:10 | SigB -10 |
| M9.1 | 116 | 43-368 | 22 | 2 | 5.6 | -75 [37] | 1.18 | 2:8 | loose |
| M18.2 | 108 | 47-355 | 6 | 6 | 0.3 | 9 [12] | 0.52 | 2:8 | RBS (GAGGTG) |
| M12.4 | 97 | 71-109 | 7 | 4 | 0.6 | -32 [3] | 3.87 | 10:10 | SigB -35 |
| M69.4 | 87 | 59-128 | 15 | 8 | **10.9** | -30 [53] | 1.75 | 10:10 | CcpA (CRE-box) |
| M42.8 | 62 | 14-295 | 17 | 0 | 0.2 | -86 [21] | 0.61 | 2:4 | loose |
| M31.4 | 39 | 16-81 | 23 | 8 | **18.8** | -40 [61] | 1.88 | 2:2 | Rex |
| M68.1 | 31 | 20-47 | 20 | 5 | **15** | -46 [25] | 3.00 | 10:10 | LiaR |
| M58.7 | 27 | 14-40 | 19 | 6 | **9.9** | -47 [56] | 3.23 | 9:10 | - |
| M62.3 | 26 | 18-34 | 20 | 13 | **16.2** | -16 [67] | 1.89 | 10:10 | Fur |
| M38.10 | 22 | 18-38 | 15 | 8 | **12.1** | -19 [28] | 2.36 | 9:10 | LexA |
| M53.9 | 20 | 8-50 | 20 | 8 | **15.1** | -47 [60] | 1.52 | 10:10 | VirR |
| M2.6 | 19 | 12-53 | 9 | 4 | 1.1 | -49 [2] | 1.12 | 3:5 | Spx |
| M13.7 | 19 | 7-48 | 21 | 5 | 1.4 | -58 [69] | 0.89 | 2:3 | - |
| M61.6 | 14 | 9-16 | 25 | 14 | **16.3** | -29 [3] | 0.81 | 9:10 | BglR2 |
| M54.1 | 14 | 6-40 | 23 | 8 | 1 | -56 [61] | 1.04 | 2:5 | - |
| M61.4 | 13 | 7-25 | 25 | 6 | 0.9 | -37 [62] | 1.20 | 2:2 | - |
| M50.10 | 13 | 7-43 | 21 | 5 | **9.6** | -77 [28] | 0.93 | 2:2 | - |
| M70.6 | 11 | 7-19 | 24 | 22 | **19.2** | -44 [63] | 1.28 | 10:10 | - |
| M3.1 | 9 | 4-20 | 21 | 9 | **15.5** | -43 [62] | 1.52 | 10:10 | - |
| M33.8 | 8 | 7-13 | 22 | 11 | 4.2 | -24 [23] | 2.78 | 9:10 | SigL |
| M49.4 | 7 | 6-11 | 23 | 12 | **16.8** | -35 [19] | 10.55 | 10:10 | PrfA |
| M20.1 | 7 | 5-7 | 25 | 16 | 5.4 | -55 [3] | 6.64 | 10:10 | - |
| M17.4 | 7 | 5-14 | 24 | 9 | **14.8** | -39 [61] | 1.14 | 3:4 | CcpB |
| M73.4 | 6 | 4-7 | 20 | 5 | 4.2 | -44 [8] | 5.03 | 6:6 | - |
| M61.3 | 6 | 5-10 | 25 | 22 | 6.4 | -33 [4] | 3.14 | 10:10 | - |
| M2.1 | 6 | 4-8 | 22 | 9 | **13** | -33 [56] | 1.05 | 2:5 | - |
| M18.6 | 4 | 3-8 | 25 | 9 | **17.2** | -51 [55] | 1.25 | 6:7 | - |
| M29.1 | 3 | 2-6 | 25 | 1 | 3 | -39 [62] | 1.64 | 9:10 | loose |
| M59.2 | 2 | 0-13 | 23 | 2 | 5.1 | -52 [63] | 2.43 | 4:5 | loose |

Fig 4 shows several examples of estimated links between expression covariates and motif occurrences as captured by the extended probit model, including covariates that were binarised or not and trees. The average number of expression covariates used (i.e. with *t*_*m,c*_ = 1) to describe the patterns of presence/absence of each of the 40 motifs in the set of sequences ranged from 1.02 (for M61.5) to 6.45 (for M71.2). Distinguishing the two types of covariates, the range was 0.28 to 4.94 for the covariates of type “vector” and 0.60 to 2.65 for the covariates of type “tree”. Fig 4ABC show three examples (M33.8, M20.1, and M68.1) of strong associations defined as those where a covariate has an estimated posterior probability of being used greater than 0.9. As expected given that we did not try to avoid redundancy between the covariates, strong association with a specific covariate did not cover all the cases of clear association between the presence/absence of a motif and the expression covariates (Fig 4DEF). Nevertheless a total of 9 motifs were strongly associated with one covariate of type vector and remarkably all these associations concerned covariates defined by ICA (none defined by PCA).

**Figure 4:**
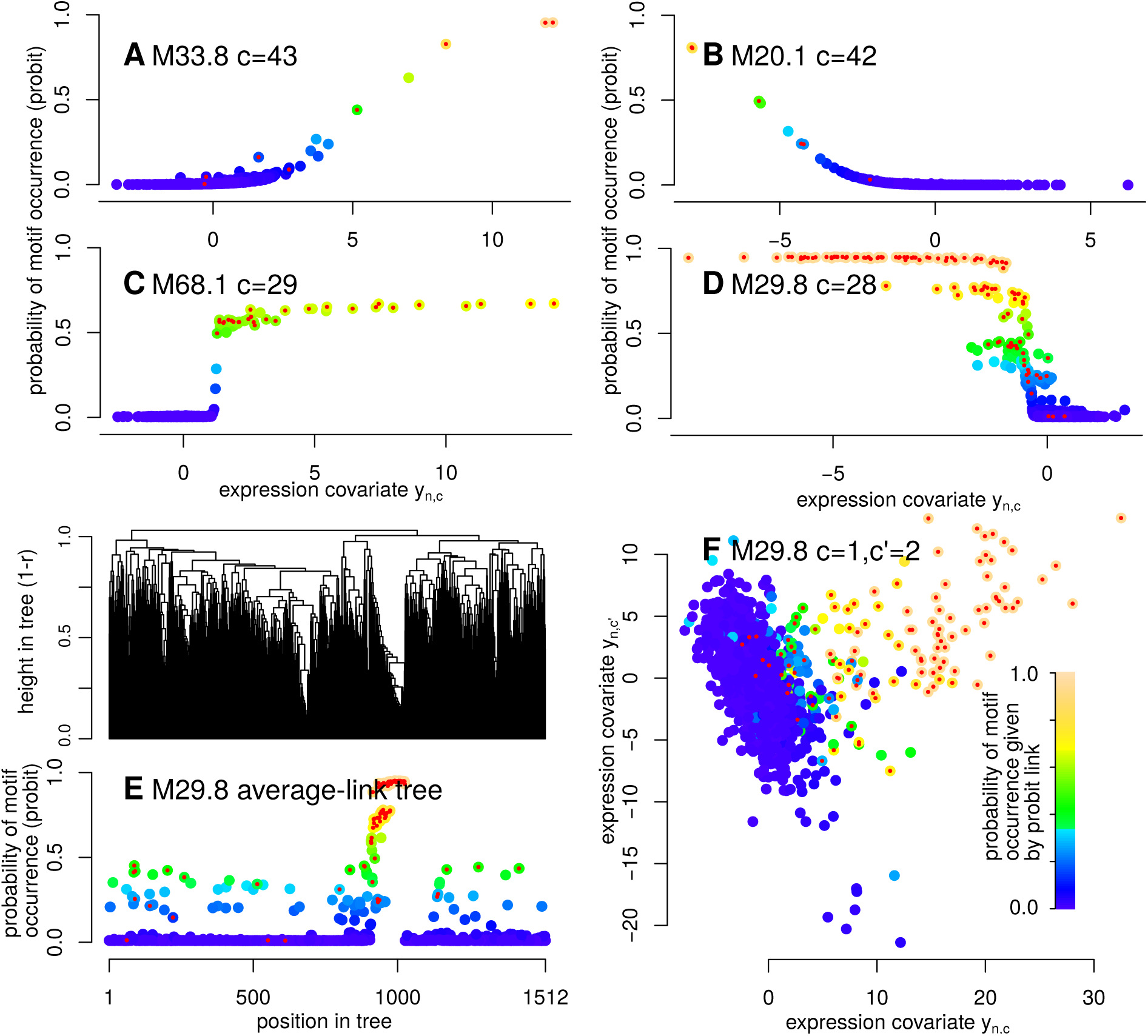
Estimated links between expression covariates and motif occurrences. Examples are shown for different motifs and covariates (indicated in subplot titles). Colour scale from blue to yellow reflects the estimated probability of motif occurrence given by the extended probit model (i.e. summarising the information from the expression data). Red dots indicate sequences in which the motif is found (estimated posterior probability of motif occurrence was ≥ 0.5).

### From validation by comparison with known motifs and regulons to new insights on *L. monocytogenes* transcription network

The 40 distinct well supported motifs exhibit considerable diversity in terms of abundance, preferred position with respect to the TSS, link with expression data, and PWM characteristics (width, information content, palindromic structure). Table 1 summarises the main characteristics of each motif. Fig 5 provides a graphical representation for two of these motifs. Similar figures are available for all motifs in S1 Fig and the numerical values of the PWM parameters are found in S1 File. Lists of genes associated with each motif and precise positions of occurrences are reported in S2 File and S3 File.

**Figure 5:**
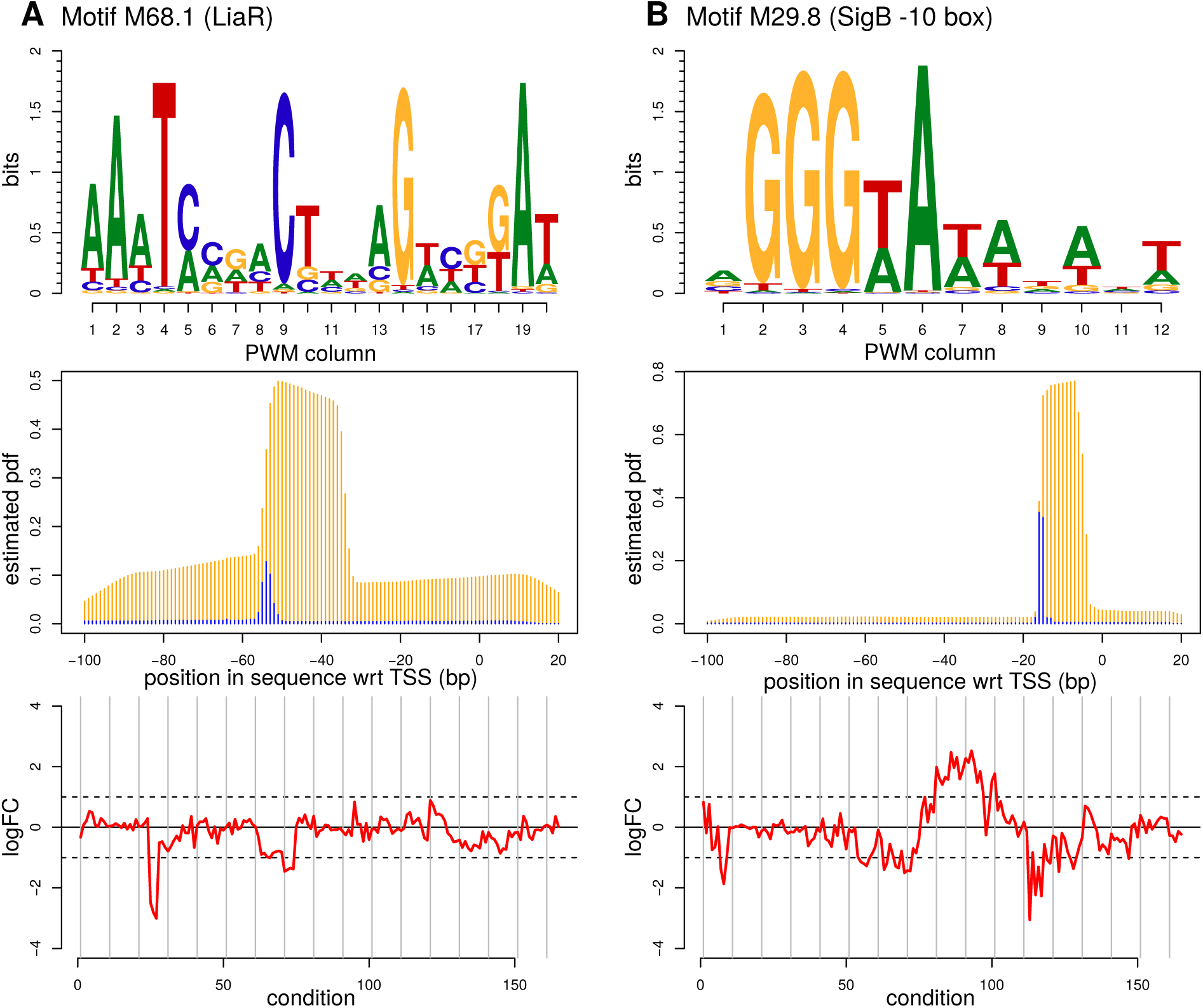
Two examples of motifs for known TFs rediscovered by our algorithm. These motifs were identified as corresponding to the binding sites of LiaR (M68.1 subplot **A**) and as the -10 box of the SigB binding sites (M29.8 subplot **B**). First row: sequence logo. Second row: estimated probability distribution function for the 5’-end of the occurrence (in blue) and probability of having the position covered by the occurrence (in orange). These probabilities are conditional on the presence of the motif in the promoter sequence. Third row: average log fold-change of expression level for downstream genes across the 165 pairs of conditions that were used to define the expression covariates.

Three complementary approaches were used to connect the 40 motifs discovered by our *de novo* approach to known regulons. The first was comparison with lists of genes collected from tables published in several expression studies and positions of transcription factor binding sites recorded in the RegPrecise database [32]; the second approach was comparison with 188 reference PWMs derived from sequence alignments extracted from the propagated RegPrecise database (accessed in July, 2018) for different taxonomic groups in the Firmicutes phylum: *Listeriaceae* (25 PWMs) and *Staphylococcaceae* (39 PWMs) and *Bacillales* (124 PWMs). The third approach consisted in dedicated literature searches associated with a careful manual examination of (i) genes downstream the promoters in which the motif was predicted to occur (ii) conditions in which the log fold-change deviated the most from 0 (iii) characteristics of PWMs and preferred positions of motif occurrences with respect to TSSs. These lines of observation provided convergent clues for many of the identified TFs.

Connections to known regulons are reported in the rightmost column of Table 1. Most of the motifs with high number of occurrences were found to describe general characteristics of promoter regions (variations on the themes of SigA -10 and -35 boxes, nucleotide composition around TSS, Ribosome Binding Site). Systematic comparison with RegPrecise PWMs was particularly informative for the identification of BglR2, CcpA, CcpB, Fur, LexA, LiaR, and Rex. Among the other identified TFs, SigB and SigL were identified based on position with respect to the TSS, comparison with sigma factor consensus defined for B. *subtilis* [10], as well as overlap with previous experimental data on SigL (also known as RpoN or sigma54) and SigB regulons in *L. monocytogenes* [33, 34, 35]. S2 Fig provides a graphical representation of the prediction of the SigB regulon from the joint occurrence of boxes -10 and -35 and of the comparison with previous studies. In brief, 89 promoter regions are predicted to contain both the -10 and -35 boxes, 88 out of them were previously reported as probable members of the SigB regulon in either [34] or [35]. Spx was identified based on (i) its position with respect to the TSS just upstream of the SigA -35 box, (ii) sequence properties reported for Spx in B. subtilis described as an AGCA element at position -44 [36], and (iii) literature data on the Spx-regulon of *L. monocytogenes* [37]. PWMs found for PrfA and VirR, two key transcription regulators involved in *L. monocytogenes* virulence, were in line with previously described sequence properties [38, 39]. Taken together, correspondences with known motifs validate our approach for *de novo* discovery and suggest that other motifs listed in Table 1 may also correspond to biologically relevant motifs.

Among the predicted regulons that we have not been able to connect to literature data, the most spectacular by its size and functional homogeneity of the regulated genes (genes encoding the translation apparatus, including ribosomal proteins), is associated with motif M58.7. Importantly, even when motifs could be linked to previously known TFs, being able to identify them in a *de novo* and automated manner can shed a new light on the corresponding regulons. For instance, our results suggest that LiaR regulon may be about twice larger than previously identified by differential expression analysis of the *liaS*-deletion mutant vs. wild-type [40]. Ordered by the number of TSSs as in Table 1, the predicted regulons for which we have identified a TF are: SigB, CcpA, Rex, LiaR, Fur, LexA, VirR, Spx, BglR2, SigL, PrfA. Additional biological observations made based on these predictions are reported in S2 Appendix.

### Comparison with other methods

We compared our results to those of different algorithms able to handle the data-set of 1, 512 sequences and 1, 512 × 165 expression matrix for *de novo* motif discovery. These algorithms are based on PWM estimation from the sole sequence data-set (MEME, [6]) or search specifically for motifs connected to the expression data using motif representations that can be either consensus string written in IUPAC degenerated symbols (FIRE and RED2, [14, 5]) or PWMs (MatrixREDUCE, [16]). Input data, parameter settings, and results are detailed in S2 Appendix.

Motif discovery without auxiliary data using MEME (see S2 Appendix) returned only up to 8 motifs with E-value less than at 1. These motifs included motifs describing general properties of the *L. monocytogenes* promoter regions, such as SigA -10 box, the presence of a RBS and of T-rich elements (our M59.9), as well as more specific motifs that are relatively abundant and/or of high information content (IC1 in Table 1): M70.6, CcpA, and Fur. A variant of the sequence element described by M20.1 and M61.3 is also found.

The three algorithms (MatrixREDUCE, FIRE, and RED2) searching for motifs related to expression data returned only short motifs (lengths up to 9) due to the use of k-mers as seeds in the initial steps of the searches (see S2 Appendix). With some parameter settings, they were all able to retrieve SigB -10 box (M29.8) which presents the particularity of being abundant and to exhibit a strong link to expression profile. It is also short and contains adjacent high information content PWM columns which makes it ideally suited for approaches based on k-mers (Fig 5B). MatrixREDUCE also discovered a short motif that corresponds to the central part of PrfA. This motif presents the particularity of exhibiting, by far, the strongest link with expression data among all the motifs that we discovered (see column FC in Table 1). Thus, comparison of our results with those of these four algorithms illustrates the interest of the statistical model described in this work: it combines the benefits of the purely sequence-based approaches, like MEME, and the benefits of algorithms that can only find motifs linked to expression data like MatrixREDUCE, FIRE and RED2.

Finally, to better understand the contributions of different ingredients of our model to the results obtained on the *L. monocytogenes* data set, we examined the results of 5 sub-models that did not incorporate one or several of the three ingredients (position with respect to TSS, expression profiles, and palindromic structures). The results detailed in S2 Appendix show that all the motifs were not sensitive to the same model ingredients and many were sensitive to several ingredients which attests the interest of an integrative statistical model.

## Discussion

The model and its associated trans-dimensional MCMC algorithm provides a new integrated framework for the discovery of regulatory motifs in bacteria with the help of transcriptome data (exact positions of TSSs and expression profiles).

A novel and key point of our sequence model is to allow motif occurrence overlaps. This simplifies the model and facilitates MCMC updates when searching for multiple motifs. Importantly, we show here that it prevents different PWM components of the model to converge to the same motif, thereby providing a alternative to the heuristic consisting in searching for motifs one-at-a-time and masking predicted occurrences in subsequent searches to avoid rediscovery of the same motifs (as implemented in MEME [6]). In this aspect, explicit modelling of motif occurrence overlaps appears as a proper statistical framework to implement what [41] named “repulsion” and for which they proposed incorporating *ad-hoc* repulsive forces between parallel MCMC runs. Binding sites do overlap in bacterial genomes [42] and our choice of modelling the contribution of different motifs that overlap by averaging the nucleotide emission probability density functions (PWM columns) is the simplest but probably not the most biologically realistic. Indeed, we also considered a more complicated model in which PWM columns corresponding to motifs that overlap contribute as a function of their information content. This satisfies the intuition that if two motifs overlap and one has a strong preference for a nucleotide at a particular position whereas the other has no or little preference, the motif with the strong preference tends to “impose” its choice. In S1 Appendix, we refer to this model as the *θ*-dependent weight mixture model of motif overlap and we describe its implementation. Because of its drawbacks (a dependence structure making that *θ* can no longer be marginalised out) we have decided to use here only the simple model of motif overlap.

By modelling the sequence and incorporating expression data as covariates, the method inherits the good-behaviour of pure sequence-based approaches grounded on well-established statistical models of DNA sequences, such as implemented in MEME [6]. This can be connected to earlier works that have incorporated external data in sequence models for motif discovery via the definition of informative priors to favour motifs whose occurrences are found in regions of the genome that are more likely to contain binding sites; for instance because they are conserved between species or depleted in nucleosomes [43, 44, 45]. In our work, the relevant covariates that describe how the probability of occurrence of a motif differs between promoters are automatically selected together with the coefficients that specify their contributions and this is achieved simultaneously for many motifs. We also show how we can account for covariates with complex structures such as the positions of the sequences in a tree. The idea of using a tree whose topology and branch lengths reflect similarities between activity profiles is to provide an alternative to the use of a predefined set of co-expression clusters. It is also found in the previous model and algorithm that we specifically developed for *de novo* discovery of sigma factor binding sites [10]. However, the probabilistic models are completely different. In [10], to give to sequences which are close in the tree (hence in the expression space) a greater chance to harbour binding sites for the same sigma factor, the occurrences of the different possible motifs are modelled as resulting from an “evolution” process along the branches of the tree. The motivation for introducing here the use of a probit model was to handle more complex representations of the expression space than a single tree. With our extended probit model, several concurrent description of the expression space can coexist (trees, continuous vectors) and the algorithm selects and combines the most relevant, with the possibility to binarise continuous vectors according to automatically adjusted cut-off values.

A limitation of our approach is that each run takes several weeks on a data-set like the one studied in this study. Most of the time is spent in the update of *a* (positions of the motif occurrences). Thus, time complexity of a MCMC sweep is approximately proportional to the total length of the sequences times the total width of the PWMs plus a small term roughly in the square of the total width for taking motif occurrence overlaps into account. Furthermore, the analysis that we conducted is based on several runs that serve to unambiguously identify peaks in the posterior landscape. These peaks are identified by the convergence of several runs to the same neighbourhood in a space of very large dimension. Here, we limited our analysis to 10 runs that were conducted in parallel. These 10 runs allowed to identify 40 stable motifs but it would not be surprising to find several other motifs by adding more runs, since some of the biologically known motifs have here only be found by 2 or 3 runs (Rex, Spx, CcpB). Future studies on other data-sets may include more runs. Computational cost is however compensated by simultaneous search for all motifs using all the expression data.

As mentioned in the introduction, *L. monocytogenes* was well-suited for a proof-of-principle application of the method. Its regulatory network contains features shared with the related Gram-positive model bacterium *B. subtilis* and features that are specific, such as those involved in pathogenicity. Comparison of our list of 40 motifs with literature data validated the method by proving its ability to re-discover, in a pure *de novo* manner, regulons of many known TFs.

In essence, the method is based on the search for “over-represented” motifs whose modelling significantly improves the probabilistic representation of the DNA sequence as measured in the likelihood. It is thus only adapted to identify the regulons of TFs playing the coordinating roles of regulating the expression of several to many transcription units. These so-called global transcription factors are opposed to local TFs that account for the vast majority of TFs in bacteria but regulate only one or very few targets in a specific biological pathway [46, 47]. For regulons of TFs that have been previously subjected to analyses by transcriptomics (e.g. SigB, LexA, Fur, LiaR, PrfA, VirR), our results contain *de novo* predictions based on the presence of motif occurrences made in a unified framework incorporating experimentally determined TSS position and expression data. This is interesting since contributions of direct and indirect regulations have not always been fully disentangled in literature by identification of transcription factors binding sites. Our results also contain the first published global predictions for the regulons of several TFs whose importance is suggested by knowledge in other bacteria such as *B. subtilis*, but which have not yet been experimentally studied in *L. monocytogenes* (CcpA, Rex, and Spx). Detection of Spx binding sites is a particularly striking achievement since its consists of a very short motif with a constrained position of occurrence directly upstream of SigA -35 box which remained elusive until the use of dedicated ChIP experiments and regression analyses in *B. subtilis* [36].

A motivation of our work was to allow the discovery of new important regulons. An interesting result, is the identification of a partially palindromic motif whose central PWM columns corresponds to palindromic consensus `ACGTAYYCGT` (M58.7 in Table 1 and S1 Fig) whose occurrences were consistently found upstream of genes encoding the translation apparatus suggesting a role in the control of growth rate. In the Gram-positive model bacterium *Bacillus subtilis*, as well as in *Escherichia coli*, by a mechanism known as the stringent response [48, 49]. In both bacteria, and probably also in *L. monocytogenes* [50], this regulatory mechanism acts by decreasing the production of the translation apparatus components when the resources in the medium become too scarce. Univocal coupling between available resources and growth rate is however not necessarily the most appropriate in all circumstances. Existence of dedicated regulatory mechanisms allowing to keep control of the growth rate, even in the presence of nutrients, may thus not seem unexpected, in particular for an intracellular pathogen like *L. monocytogenes*. We hypothesise that the new regulon that we detected in *L. monocytogenes* may be an instance of such a mechanism whose biological role and regulatory molecules remain to be identified.

## Supporting information

Detailed description of the statistical model and the associated MCMC algorithm.

## Supporting information

**S1 Fig. Summary plots for the 40 stable motifs.**

**S2 Fig. Prediction of SigB regulon and comparison with previous studies. S1 File. PWMs for the 40 stable motifs.** Table in XLSX format.

**S2 File. Detailed information on the predicted regulons associated to each of the 40 stable motifs.** Table in XLSX format.

**S3 File. Most likely positions for the occurrences for the 40 motifs in the 1**,**512 promoter regions (121 bp around the selected TSSs).** Table in XLSX format.

**S1 Appendix. Detailed description of the statistical model and the associated MCMC algorithm.**

**S2 Appendix. Biological observations, comparison with results of some other algorithms and contribution of different sources of information to motif discovery.**

## Data accessibility

The source code of `Multiple` is distributed under the GNU General Public License (https://forgemia.inra.fr/pierre.nicolas/multiple); expression covariates and sequence data used in this study are made available with the program. Supplementary materials provide a full description of the methods and results.

## Authors’ contributions

I.S., S.S., V.F. and P.N. designed the study, interpreted the results, contributed to the writing of the manuscript, and gave final approval for publication. I.S. and P.N. implemented the algorithm and processed the data.

## Acknowledgements

We thank Birgitte H. Kallipolitis, Pascal Piveteau, and the List Maps consortium for helpful discussions; INRA MIGALE bioinformatics facility (doi: 10.15454/1.5572390655343293E12) for providing computational resources.

