## Supplementary material for "Statistical modelling of bacterial promoter sequences for regulatory motif discovery with the help of transcriptome data: application to *Listeria monocytogenes*": Detailed description of the statistical model and the associated MCMC algorithm.

Ibrahim Sultan[1], Vincent Fromion[1], Sophie Schbath[1], Pierre Nicolas[1]
**1** MaIAGE UR1404, INRA, Université Paris-Saclay, Jouy-en-Josas, France

October 29, 2019

**References** **43**

This appendix is divided into two sections: Model (p. 4) and MCMC algorithm (p. 20). As needed to provide here a comprehensive description of the model, the first section is partially redundant with the main text of manuscript.

### 1 Model

We introduce in this section the assumptions of our probabilistic model and the notations used for the data, hidden variables, and parameters. The Directed Acyclic Graph (DAG) represented in Figure 1 provides an overview of the model and can be used as a summary of the notations (more details on the DAG are found in 1.3.3).

#### 1.1 Sequence model

##### 1.1.1 Probabilistic modelling of the sequences

The sequence data set that we consider is composed of a number $\mathcal{N}$ of DNA sequences, each of the same length $\mathcal{L}$. Let denote $x = (x_n)_{n=1:\mathcal{N}}$ the set of DNA sequences and $x_{n,l}$ the nucleotide in position $l$ of the sequence of index $n$ [1] ($x_{n,l} \in \{\mathtt{A}, \mathtt{C}, \mathtt{G}, \mathtt{T}\}$). Our probabilistic representation for these sequences involves $\mathcal{M} + 1$ unobserved components that capture the heterogeneity of the nucleotide composition along the sequences. These components consists of $\mathcal{M}$ motif models with respective widths $(w_1, ...., w_{\mathcal{M}})$ and one background model.

The description of the probabilistic model for the sequences $x$ contains two aspects: the modelling of positions of motif occurrences and the modelling of the sequence given the positions of these occurrences. The modelling of the positions of motif occurrences proceeds as follows.

- The model assumes zero or one occurrence of each of the $\mathcal{M}$ motifs per sequence. In a sequence, the occurrences of the $\mathcal{M}$ motifs are furthermore modelled as mutually independent.

- The probability that an occurrence of motif $m$ occurs in sequence $n$ is denoted $\alpha_{m,n}$ with

$$\alpha_{m,n} = \Phi(\beta_0 + \sum_c \mathbb{I}\{t_{m,c} = 1\}\beta_{m,c}\tilde{y}_{m,n,c}) \qquad (1)$$

  where $\Phi$ is the cumulative distribution function of the standard Gaussian and $\tilde{y}_{m,n,c}$ is a function of $y_{n,c}$, $g_m$, and $h_m$ (see Equation 11 and 12). The focus of section 1.2 is this modelling to incorporate expression data in $\alpha_{m,n}$, thereby bringing an information which is specific of each sequence.

[1] We will use the terms “sequence $n$” to refer to the” sequence of index $n$” and the same for the other components of the model (motifs, regions, . . .).

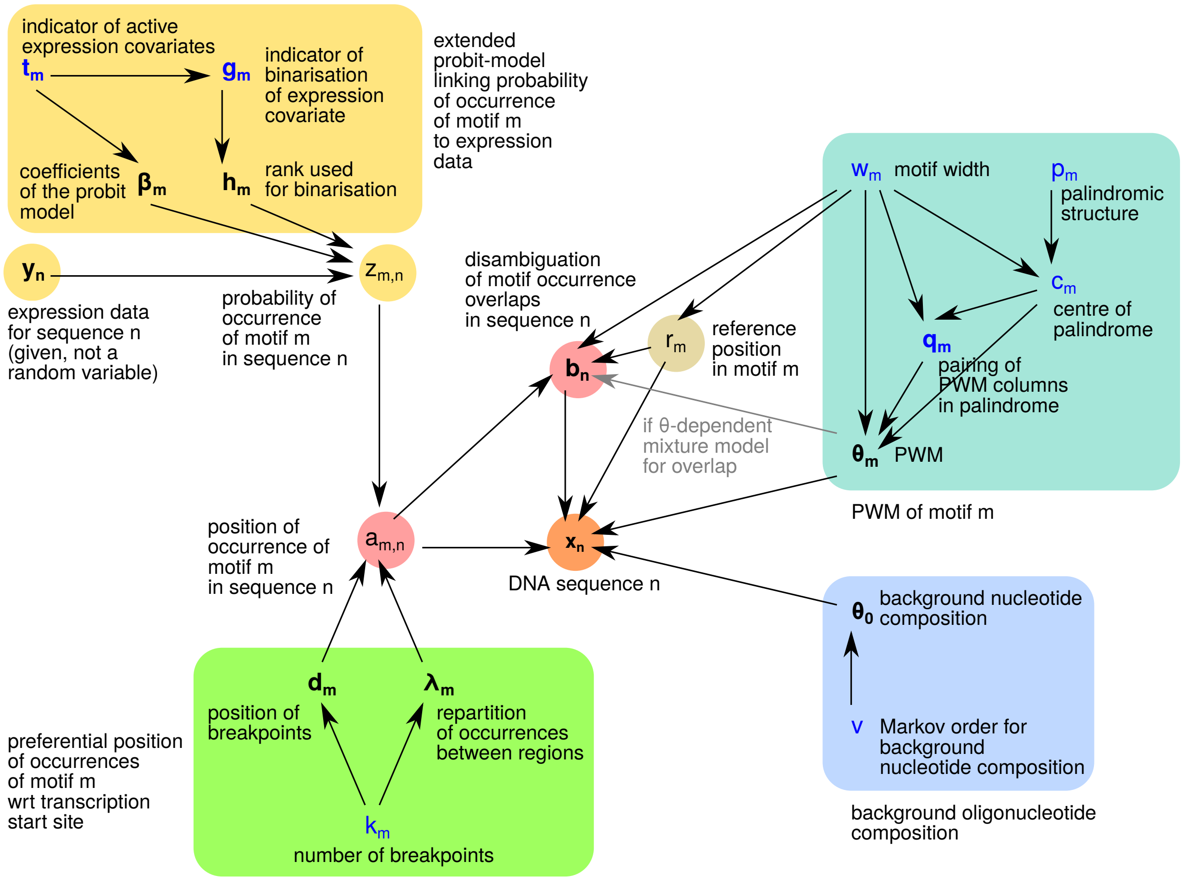

Figure 1: DAG of our model. Vertices represent the variables (parameters, hidden variables, observed data) and edges show the factorisation of the joint probability distribution (see Equation 34). Coloured areas highlight groups of variables that contribute to a same aspect of the model. The upper-left part of the DAG dedicated to the incorporation of expression data in the sequence model which is presented in section 1.2. In this representation we show the variables for only one motif $m$ and one sequence $n$. The grey edge that links the disambiguation hidden variable $B$ to $\theta$ is specific to the model of overlap that takes into account information content of the columns of the PWM ($\theta$-dependent mixture). Of note, our implementation of the MCMC algorithms makes this option incompatible with the modelling of palindromic structures via the variables $(p_m, c_m, q_m)$. Multidimensional variables (vectors or matrices) are boldfaced; variables whose values change the dimension of the model are in blue; continuous random variables are denoted by Greek letters.

- The probability distribution for the position of motif $m$ in a sequence is defined by a piecewise constant function with $k_m$ breakpoints. The positions of the breakpoints are denoted $d_m = (d_{m,k})_{k=1:k_m}$, where $1 < d_{m,k} < d_{m,k+1} \leq \mathcal{L}$. The probability of occurrence of the motif at a particular position $l$ in region $k$ (i.e. $d_{m,k-1} \leq l < d_{m,k}$) is $\lambda_{m,k}/(d_{m,k} - d_{m,k-1})$ where $d_{m,k} - d_{m,k-1}$ corresponds to the length of region $k$. Hence, for a given number of occurrences of motif $m$, the distribution of these occurrences between the $k_m + 1$ regions is described by a multinomial of parameters $\lambda_m = (\lambda_{m,k})_{k=1:k_m+1}$, with $\lambda_{m,k} \geq 0$ and $\sum_{k=1}^{k_m+1} \lambda_{m,k} = 1$, where $\lambda_{m,k}$ is the probability for an occurrence of motif $m$ to be found in the region $k$.

- For practical reasons that pertain to the implementation of the update of the motif width $w_m$ in the MCMC algorithm we do not always rely on the first position within motif $m$ to index its position. Instead, we use a reference position $r_m \in \{1, \ldots, w_m\}$ that will allow the motif occurrences to overlap the sequence boundaries. According to this modelling, a motif can extend up to $r_m - 1$ positions upstream of the sequence (when its indexed position occurs at position 1 in the sequence) and up to $w_m - r_m$ positions downstream of the sequence (when its indexed position occurs at position $\mathcal{L}$ in the sequence).

The modelling of the nucleotide sequence given the positions of motif occurrences involves a set of parameter for each model component: $(\theta_1, ...., \theta_{\mathcal{M}})$ for the $\mathcal{M}$ motifs and $\theta_0$ for the background.

- In keeping with the simple PWM framework for motif modelling, the nucleotide at position $w \in \{1, \ldots, w_m\}$ inside an occurrence of motif $m$ is drawn from a multinomial distribution of parameters $\theta_{m,w} = (\theta_{m,w,u})_{u\in\{\mathtt{A,C,G,T}\}}$, with $\theta_{m,w,u} \in [0, 1]$ and $\sum_u \theta_{m,w,u} = 1$, where $\theta_{m,w,u}$ is the probability to observe nucleotide $u$ in the position $w$ of the motif $m$. As explained below the columns inside a PWM are not necessarily modelled with $w_m$ independent vectors $\theta_{m,w}$'s since we account for the possibility of (fully or partially) palindromic motifs (see page 9).

- Nucleotides outside motif occurrences are modelled as generated by a background model. The background model that we choose is a homogeneous Markov model of order $v \in \{0, 1, \ldots\}$. We denote $\theta_0 = ((\theta_{0,s,u})_{s\in\{\mathtt{A,C,G,T}\}^{v'}, u\in\{\mathtt{A,C,G,T}\}})_{v'=0:v}$ the parameters of this model, where $\theta_{0,s,u}$ corresponds to the probability of nucleotide $u$ after the word $s$ of length $v$. This model involves $4^v \times 3$ independent parameters (since $\sum_{u\in\{\mathtt{A,C,G,T}\}} \theta_{0,s,u} = 1$) that allow to account for the composition in word of length $v+1$; $\theta_0$ also encompasses the transitions matrices of order $v'$ from 0 to $v - 1$ that serve to model the nucleotide sequences at position $l \leq v$ (i.e. the initial distribution of the Markov model).

- The assumption of independence between the occurrences of the $\mathcal{M}$ motifs allows overlaps. Two modelling options were envisioned to model nucleotide composition at positions where two or more motif occurrences overlap.
  - The simplest model consisted of considering a simple arithmetic mean of the probability density functions associated with the different motifs that overlap. We refer to this model as the equal weight mixture model for motif overlaps.
  - A more general model was also implemented and consisted in a weighted mean of the probability density functions associated with the different motifs. We refer to this model as a $\theta$-dependent weight mixture model for motif overlaps. According to this model, each density function described by $\theta_{m,w}$ receives a weight that increases with the level of constraint that it imposes on the nucleotide to be generated. We considered for this purpose a quantity denoted $IC(\theta_{m,w})$ expressed in information bits and often referred to as the 'information content' of a column of a PWM in the literature on biological sequence motifs. $IC(\theta_{m,w})$ takes values from 0 when the randomness is maximum (i.e. uniform distribution) to 2 in absence of randomness (i.e. $\theta_{m,w,u} = 1$ for one of the possible nucleotide $u \in \{\mathtt{A}, \mathtt{C}, \mathtt{G}, \mathtt{T}\}$). It is computed as

    $$IC(\theta_{m,w}) \;=\; 2 - \sum_{u \in \{\mathtt{A},\mathtt{C},\mathtt{G},\mathtt{T}\}} \theta_{m,w,u} \log_2 \theta_{m,w,u} \,, \qquad (2)$$

    where the sum term corresponds to the Shannon entropy of the nucleotide distribution described by $\theta_{m,w}$.

##### 1.1.2 Hidden variables

In order to be able to work with this model, we introduced two layers of hidden variables which are:

- $A = (A_{m,n})_{m=1:\mathcal{M},\, n=1:\mathcal{N}}$ is a layer of random variables that encode the position of the occurrences of the motifs, with $A_{m,n} \in \{0, 1, \ldots, \mathcal{L}\}$ being the position of motif $m$ in sequence $n$ (0 encodes the absence of occurrence).

- $B = (B_{n,l})_{n=1:\mathcal{N},\, l=1:\mathcal{L}}$ is a layer of random variables that encode the disambiguation of the overlaps between motifs, with $B_{n,l} \in \{0, 1, \ldots, \mathcal{M}\}$ denoting the model component (background or one of the $\mathcal{M}$ motifs) responsible for the probability distribution function of the nucleotide $x_{m,n,l}$. This idea of disambiguation by a dedicated hidden variable is made possible by our choice of modelling the nucleotide emission probability at positions where motifs overlap as a mean since the resulting density can be seen as the marginal of a mixture model (equal weight mixture model or $\theta$-dependent weight mixture model).

Having introduced these hidden random variables $(A, B)$ allows to define the complete data $(x, A = a, B = b)$ whose density function given $y$ and the parameters

$$(\theta_0, \theta_{1:\mathcal{M}}, t_{1:\mathcal{M}}, \beta_{1:\mathcal{M}}, g_{1:\mathcal{M}}, h_{1:\mathcal{M}}, d_{1:\mathcal{M}}, \lambda_{1:\mathcal{M}})$$

that we will simply write $(\theta, t, \beta, g, h, d, \lambda)$ decomposes as

$$\begin{aligned} \pi(x, a, b|\theta, y, t, \beta, g, h, d, \lambda) &= \prod_{n=1:\mathcal{N}} \pi(x_n, b_n, a_n|\theta, y, t, \beta, g, h, d, \lambda) \\ &= \prod_{n=1:\mathcal{N}} \pi(x_n|b_n, a_n, \theta)\pi(b_n|a_n, \theta)\pi(a_n|y, t, \beta, g, h, d, \lambda)\,, \end{aligned} \tag{3}$$

where the terms $\pi(x_n|b_n, a_n, \theta)$, $\pi(b_n|a_n, \theta)$, $\pi(a_n|y, t, \beta, g, h, d, \lambda)$ are easy to compute as shown below.

The density function of the sequence given the hidden variables writes

$$\begin{aligned} \pi(x_n|b_n, a_n, \theta) &= \prod_{l=1:\mathcal{L}} \pi(x_{n,l}|b_{n,l}, a_n, \theta) \\ &= \prod_{l=1:\mathcal{L}} [\theta_{b_{n,l}, l-a_{n,b_{n,l}}+1, x_{n,l}}]^{\mathbb{I}\{b_{n,l}\geq 1\}} [\theta_{0, x_{n,l-v:l-1}, x_{n,l}}]^{\mathbb{I}\{b_{n,l}=0\}}\,, \end{aligned} \tag{4}$$

where $\mathbb{I}\{z\}$ is the indicator function that takes value 1 if the Boolean variable $z$ is true and 0 otherwise, and $x_{n,l-v:l-1}$ denotes the word of length $v$ finishing at position $l-1$ of sequence $n$, i.e. $x_{n,l-v:l-1} = (x_{n,l-v}, \ldots, x_{n,l-1})$.

The density function of the disambiguation hidden variables given the position of the motif occurrences writes

$$\pi(b_n|a_n, \theta) = \prod_{l=1:\mathcal{L}} \pi(b_{n,l}|a_n, \theta)\,, \tag{5}$$

where to write $\pi(b_{n,l}|a_n, \theta)$ it is easier to introduce a variable $o_{n,l}$ corresponding to the number of motif occurrences that overlap the position $(n, l)$, we define

$$o_{n,l} = \sum_{m=1:\mathcal{M}} \mathbb{I}\{a_{m,n} \neq 0, a_{m,n} \leq l + r_m - 1 < a_{m,n} + w_m\}\,. \tag{6}$$

If one or several motif occurrences overlap the position $(n, l)$, we further need to distinguish the case of the equal weight mixture model for motif overlaps in which

$$\begin{aligned} \pi(B_{n,l} = m|a_n, o_{n,l}, \theta) \propto\ & \mathbb{I}\{m = 0, o_{n,l} = 0\} \\ & + \mathbb{I}\{m > 0, a_{m,n} \neq 0, a_{m,n} \leq l + r_m - 1 < a_{m,n} + w_m\}\,, \end{aligned} \tag{7}$$

and the case of the $\theta$-dependent weight mixture model for motif overlaps in which

$$\begin{aligned}
&\pi(B_{n,l} = m|a_n, o_{n,l}, \theta) \\
&\quad \propto \quad \mathbb{I}\{m = 0, o_{n,l} = 0\} \\
&\qquad +f(IC(\theta_{m,l+r_m-a_{m,n}})\mathbb{I}\{m > 0, a_{m,n} \neq 0, a_{m,n} \leq l + r_m - 1 < a_{m,n} + w_m\}\,.
\end{aligned} \tag{8}$$

The density function of the motif occurrences writes

$$\begin{aligned}
\pi(a_{m,n}|y,t,\beta,g,h,d,\lambda) \quad = \quad & \mathbb{I}\{a_{m,n} = 0\}(1 - \alpha_{m,n}) \\
& +\mathbb{I}\{a_{m,n} \neq 0\}\alpha_{m,n} \prod_{k=1:k_m+1} \left[\frac{\lambda_{m,k}}{d_{m,k} - d_{m,k-1}}\right]^{\mathbb{I}\{d_{m,k-1} \leq a_{m,n} < d_{m,k}\}},
\end{aligned} \tag{9}$$

where $\alpha_{m,n}$ is given by Equation 1 and depends is a function of $y, t, \beta, g, h$.

###### 1.1.3 Dimension and palindromic constraints

The dimension of the model expressed as the number of parameters that serve for the probabilistic description of the sequence data are not fixed and will be adjusted in the course of the estimation algorithm. In particular, the dimension of the model change with:

- the number of breakpoints used to describe the distribution of the occurrences of the motifs $(k_m)_{m=1:\mathcal{M}}$,
- the widths of the $m$ motifs $(w_m)_{m=1:\mathcal{M}}$,
- the Markov order $v$ of the background,
- the palindromic constraints that are applied to the PWMs.

Many of the known binding motifs of TFs are palindromic in the sense that pairs of columns at opposed positions with respect to the centre of the motif appeared as mirrored according to Watson-Crick base pairing rule (`A` : `T` and `C` : `G`). We considered that modelling these palindromic constraints on PWMs could be useful since they reduce the dimension of the model and therefore simultaneously increase the amount of data available to estimate each parameter and decrease the size of the search space for the parameter values.

Instead of imposing a strict palindromic constraint on all or a subset of the motifs we developed a more flexible modelling approach that allows, for each motif $m$, smooth transitions between palindromic and non palindromic structures. This approach relied on the introduction of the following variables to define the active constraints on $\theta_m$:

- $p_m$, a binary variable taking value 1 if the motif $m$ is allowed to have a (partially) palindromic structure, 0 otherwise;

- $c_m \in \{1.5, 2, 2.5, \ldots, w_m - 0.5\}$, a discrete variable used when $p_m = 1$, to record the position of the centre of the palindromic structure; the range for $c_m$ allows two types of palindromic structure ("even" and "odd" types, where the odd type contains a central unpaired column at $w = c_m$);

- $q_{m,w}$, a binary variable used when $w \geq 2c_m - w_m$ and $w < c_m$ to indicate whether columns $w$ and $2c_m - w$ are paired, i.e. $\theta_{m,w,u} = \theta_{m,2c_m-w,\bar{u}}$ when $(r, \bar{u})$ is a Watson-Crick pair.

The motivations for these modelling choices that allow intermediate levels of constraints between the non palindromic and the fully palindromic structures stem from several considerations. First, our overarching goal was to set up an algorithm that could identify as many motifs as possible simultaneously which is incompatible with the idea of having all or none of the motifs palindromic. Second, allowing separate sub-populations of motif components to coexist in our model (fully palindromic and non palindromic) would have certainly caused convergence issues due to the size of the dimension jump needed to move one motif from one population to another (dividing or multiplying by two the number of free parameters in $\theta_m$). In contrast, our model that allows smooth transitions between non-palindromic and palindromic motif representation makes it possible to design algorithms that gradually increase or decrease the number of free parameters in $\theta_m$. Finally, it should also be noticed that while some level of palindromic structure are often obvious it is unclear to which extent the biological motifs are fully palindromic or partially palindromic.

#### 1.2 An extended probit model to use expression data as covariates

We already discussed our choice of incorporating expression data in our model as covariates that can carry information on the probability of occurrence of the motifs in the different sequences. To establish this link between expression data and probability of occurrence of a motif, we choose to adopt the methodological framework of the probit regression which presents the advantage of being relatively simple to implement, compared to the logit regression, in our context of Bayesian and MCMC-based inference. This simplicity stems from the availability of a data augmentation scheme in which a Gaussian latent variable model is introduced (Albert and Chib, 1993). We also realised that this model could easily accommodate an extension that could be seen as a finalisation of expression covariates according to an automatically adjusted breakpoint which is very appealing in our modelling context for two reasons. First, it allows to model sharp switches in the probability of occurrences as a function of the position of the sequence in the expression space without imposing the probability of occurrence to jump between 0 and 1 between the sides of the breaks. Second, this binary representation of the covariate makes it possible to incorporate whole tree structures in the regression model. In this case, the finalisation breakpoint is located along the branches of the tree instead of along a simple axis. In this section we start by describing our model for incorporation of covariates that have the form of a vector of continuous variables. Then, we describe (page 12) how the version of the model in which the covariate is binaries can also handle trees.

##### 1.2.1 Vectors of continuous variables

We consider the case of a number $\mathcal{C}$ of vectors of continuous variables denoted $y = (y_{n,c})_{n=1:\mathcal{N},c=1:\mathcal{C}}$ that are to be used as covariates in the modelling of the events $\mathbb{I}\{A_{m,n} > 0\}$ (i.e. an occurrence of motif $m$ is found in sequence $n$) for $n = 1 : \mathcal{N}$. Each motif $m$ is modelled independently. According to the standard probit model, we would write,

$$\pi(A_{m,n} > 0|y) \quad = \quad \Phi(\beta_{m,0} + \sum_{c=1:\mathcal{C}} \beta_{m,c} y_{n,c}) \,, \tag{10}$$

where the $\beta$'s are the regression coefficients ($\beta_{m,c} \in \mathbb{R}$) and $\Phi$ is the cumulative distribution function of the standard normal distribution which serves to map any value of $\beta_{m,0} + \sum_{c=1:\mathcal{C}} \beta_{m,c} y_{n,c}$ into a probability between 0 and 1.

To allow the automatic selection of the covariates that are relevant to predict the occurrences of motif $m$ (choice of model dimension) and the aforementioned binarisation of the covariates, our model involves the following parameters (variables in our Bayesian inference context),

- $t_m = (t_{m,1}, \ldots, t_{m,C})$ where $t_{m,c} \in \{0, 1\}$ indicates whether covariate $c$ should be taken into account in the probability of occurrence of motif $m$ (i.e. $A_{m,n} > 0$);
- $\beta_m = (\beta_{m,0}, \beta_{m,1}, \ldots, \beta_{m,C})$ where $\beta_{m,c} \in \mathbb{R}$ represents when $t_{m,c} = 1$ the parameter used in a probit regression that relates $y_{n,c}$ to the probability that $A_{m,n} > 0$, $\beta_{m,0}$ is the intercept parameter;
- $g_m = (g_{m,1}, \ldots, g_{m,C})$ where $g_{m,c} \in \{0, 1\}$ indicates, when $t_{m,c} = 1$, whether the values in the vector $y_c$ are binarised ($g_{m,c} = 1$) or enter directly as they are in the probit model ($g_{m,c} = 0$);
- $h_m = (h_{m,1}, \ldots, h_{m,C})$ where $h_{m,c} \in \{1, \ldots, \mathcal{N} - 1\}$ indicates, when $t_{m,c} = 1$ and $g_{m,c} = 1$, the rank in the vector $y_c$ of the value used for binarisation; we use the notation $y_{(h_{m,c}),c}$ for the corresponding cut-off.

In keeping with the probit regression framework, we write the probability of occurrence of motif $m$ in sequence $n$ as

$$\pi(A_{m,n} > 0|y, t, \beta, b, h) \quad = \quad \Phi\big(\beta_{m,0} + \sum_{c} \beta_{m,c} \mathbb{I}\{t_{m,c} = 1\} \tilde{y}_{m,n,c}\big) \,, \tag{11}$$

where $\tilde{y}_{m,n,c}$ corresponds to $y_{n,c}$ after an eventual motif-specific binarisation and writes

$$\begin{aligned}\tilde{y}_{m,n,c} &= \mathbb{I}\{g_{m,c}=0\}y_{n,c} \\ &\quad +\mathbb{I}\{g_{m,c}=1\}\left[\Big(\frac{h_{m,c}}{\mathcal{N}}-1\Big)\mathbb{I}\{y_{n,c}\leq y_{[h_{m,c}],c}\}+\frac{h_{m,c}}{\mathcal{N}}\mathbb{I}\{y_{n,c}>y_{[h_{m,c}],c}\}\right].\end{aligned} \quad (12)$$

When binarisation is active (case $g_{m,c}=1$), this formula maps $y_{n,c} > y_{[h_{m,c}],c}$ to $h_{m,c}/\mathcal{N}$ and $y_{n,c} \leq y_{[h_{m,c}],c}$ to $h_{m,c}/\mathcal{N}-1$. Any mapping to other values than $h_{m,c}/\mathcal{N}$ and $h_{m,c}/\mathcal{N}-1$ would be equivalent in terms of the distribution of $\mathbb{I}\{A_{m,n}>0\}$ that it allows, provided that the $\beta$ are readjusted. However, this mapping was chosen because it presents a two-fold advantage: it ensures the centring to 0 of $\tilde{y}_{m,n,c}$ whatever the value of $h_{m,c}$ when $g_{m,c}=1$ and it ensures a difference of 1 between the two possible values of the binarised covariates (hence preserving the interpretation of $\beta$ across the possible values for $h_{m,c}$). The centring is important for the mixing of the MCMC algorithm since the $\mathcal{C}$ groups of variables $(t_{m,c}, g_{m,c}, \beta_{m,c}, h_{m,c})$ are updated successively each $c$ (see algorithm overview in section 2.1.2).

Figures 2 and 3 intend to illustrate the shapes of the relationship between expression covariates and the probability of motif occurrence that are allowed by the simple probit model and by our extension based on covariate binarisation. In one dimension (Figure 2), the simple probit model can account for a sharp switch between a region of low probability of occurrence and a region of high probability of occurrence but these regions have then probabilities close to 0 and 1. In contrast, the extended probit model with its three parameters $h_{m,c}$, $\beta_{m,0}$ and $\beta_{m,c}$ can describe freely the position of the switch and the probability of motif occurrence on both side of switch. In two dimensions (Figure 3), the five parameters of the extended probit model can describe freely the position of the switches on each axis, but only three parameters are used to describe the probability of motif occurrence in the four regions defined by the switches. Indeed, when dimension $\mathcal{C}$ increases the number ($2^{\mathcal{C}}$) of regions delineated by the switches increases exponentially but the number $(1+2\mathcal{C})$ of parameters increases only linearly as a result of the assumption of additive effects (on the probit scale). This slow increase makes it possible to use this model even when dimension $\mathcal{C}$ is relatively high.

##### 1.2.2 Trees: branch lengths and topology

Further extending the previous model described by Equations 11 and 12, we consider that covariates can come not only in the form of vectors of continuous variables but also in the form of trees. For simplicity, we consider here rooted binary trees in which all the leaves are at a same distance from the root. In our context, these trees are obtained by hierarchical clustering of gene expression. Such a tree covariate $c$, contains a total of $2\mathcal{N}-1$ nodes that decompose into $\mathcal{N}$ terminal nodes or leaves and $\mathcal{N}-1$ internal nodes. It can be entirely encoded (topology and branch lengths) as a vector internal node heights $h_c = (h_{c,i})_{i=1:\mathcal{N}-1}$

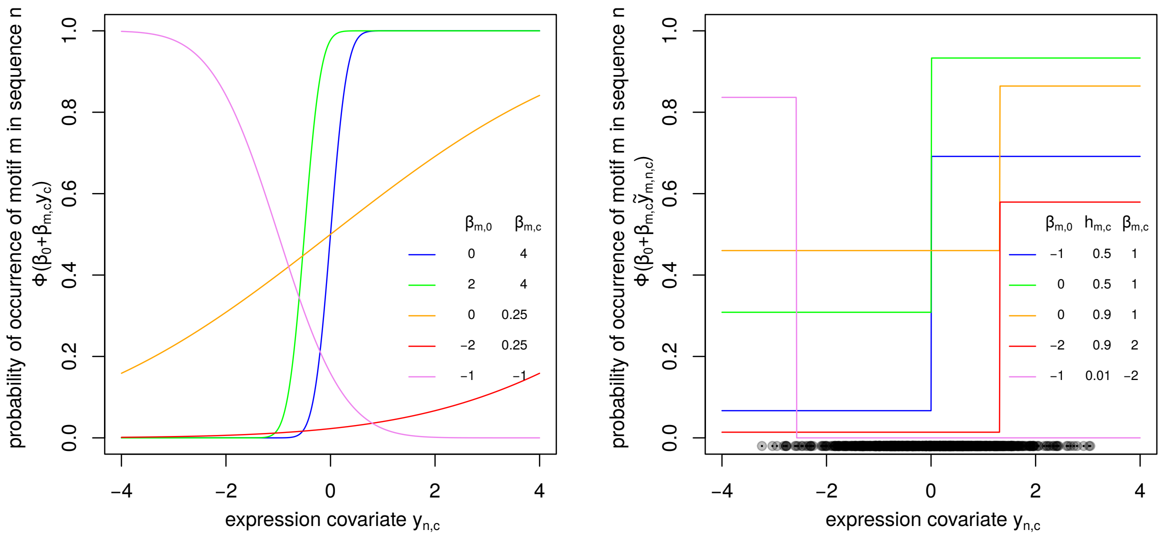

Figure 2: Graphical illustration of our probit model of probability of motif occurrence for one expression covariate. The probability of motif occurrence in sequence $n$ is represented as a function of the value of the covariate $y_{n,c}$ for five sets of parameters. Left: simple probit regression model with two parameters $\beta_{m,0}$ and $\beta_{m,c}$ that relate the value of expression covariate $c$ available for gene $n$ ($y_{n,c}$) to the probability of occurrence of motif $m$. Right: extended probit regression model binarising the expression covariate $c$ according to a cut-off parameter $h_{m,c}$ expressed as a rank (quantile instead of rank in the insert legend). The horizontally aligned grey points at the bottom of the plot represents the values of $y_{n,c}$ that served for the binarisation according to $h_{m,c}$.

ordered such as $h_{c,i} \leq h_{c,i+1}$ and a $(\mathcal{N}-1)\times 2$ matrix $s_c = (s_{c,i,1}, s_{c,i,2})_{i=1:\mathcal{N}-1}$ that identifies the subtrees merged at each internal node. In practice the $2\mathcal{N}-2$ subtrees hanging below each node are identified by values in the ranges $\{-\mathcal{N},\ldots,-1\}\cup\{1,\mathcal{N}-1\}$, with negative values corresponding to terminal nodes (in our cases the indexes of the sequences) and positive values corresponding to internal nodes ordered by height as in vector $h_c$.

Taken together, the vector $h_c$ and the two-columns matrix $s_c$ replace the vector $y_c$ when the covariate $c$ is a tree. Furthermore, the trees can be accommodated by our model only in the form of binarised covariates, hence $g_{m,c} = 1$. Compared to a covariate of type "vector", the interpretation of the $h_{m,c}$ is slightly changed such as it identifies a subtree (indexed as in the rows of matrix $s_c$). The mapping of the covariate to $\tilde{y}_{m,n,c}$, is also adapted. We used, $\tilde{y}_{m,n,c} = |h_{m,c}|/\mathcal{N}-1$ if sequence $n$ belongs to the subtree $h_{m,c}$ and $|h_{m,c}|/\mathcal{N}$ otherwise, where we use the notation $|h_{m,c}|$ for the number of sequences in the subtree $|h_{m,c}|$. Figure 4 illustrates how the selection of a subtree allows to associate different probability of motif occurrence to different sequences.

##### 1.2.3 Data augmentation for the extended probit

Following the data augmentation scheme described by Albert and Chib (1993), we introduce a unit variance Gaussian random variable $Z_{m,n}$ such that the probability modelled by the

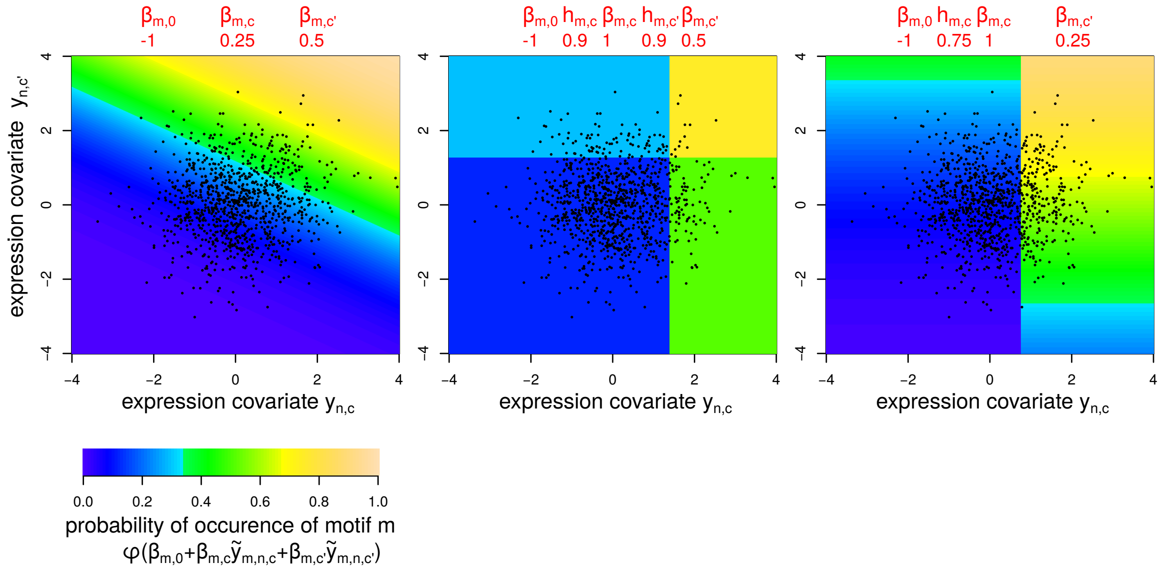

Figure 3: Graphical illustration of our probit model of probability of motif occurrence for two expression covariate. The probability of motif occurrence in sequence $n$ is represented as a function of the value of the covariates $y_{n,c}$ and $y_{n,c'}$ for three sets of parameters. Left: simple probit regression model with three parameters $\beta_{m,0}$, $\beta_{m,c}$ and $\beta_{m,c'}$ that relate the value of expression covariates $c$ and $c'$ to the probability of occurrence of motif $m$. Middle: extended probit regression model binarising the expression covariates $c$ and $c'$ according to cut-off parameters $h_{m,c}$ and $h_{m,c'}$ expressed as a rank (quantile instead of rank in the insert legend). Right: extended probit regression model binarising the expression covariates $c$ but not $c'$. The black dots represents the values of $y_{n,c}$ and $y_{n,c'}$ that served for the binarisation according to $h_{m,c}$ and $h_{m,c'}$.

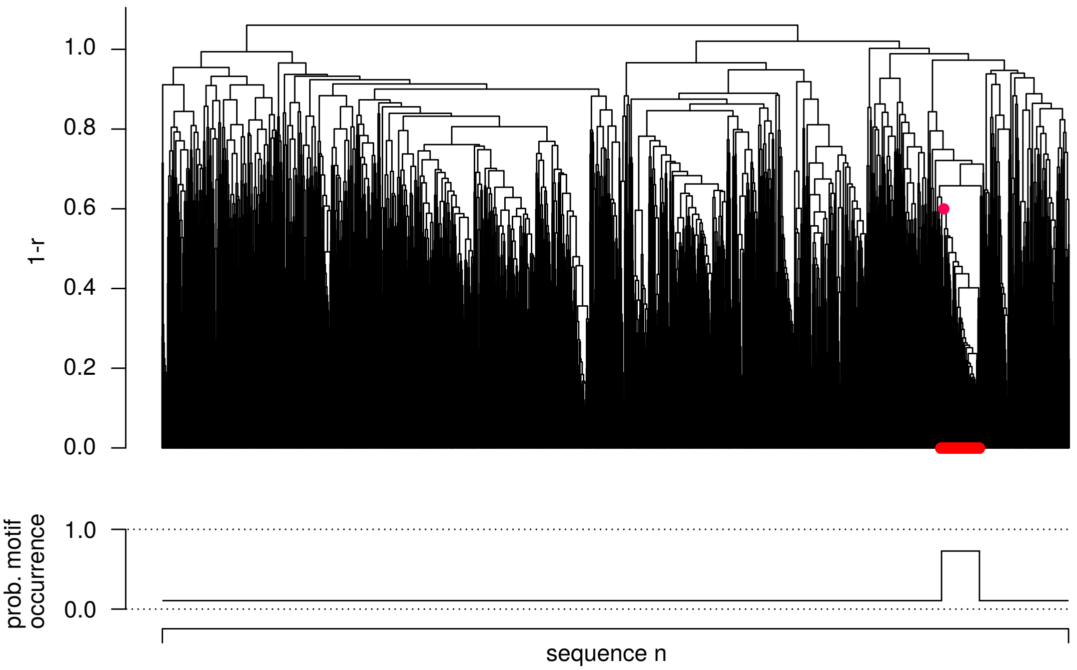

Figure 4: A graphical representation of how the process of selecting a cut on the PCA or the ICA can as well be applied on the clustering tree. In the case of the tree we are selecting a specific subtree (branch or merge) and all the sequences hanging from this subtree will have a different probability of having the motif compared to the rest of the tree.

probit regression corresponds to the probability of $Z_{m,n} > 0$. The distribution of this random variable is

$$Z_{m,n}|y, t, \beta, b, h \sim \mathcal{N}(\text{mean} = \beta_{m,0} + \sum_c \beta_{m,c}\mathbb{I}\{t_{m,c} = 1\}\tilde{y}_{m,n,c}, \text{var} = \sigma_z^2)\,, \tag{13}$$

where $\sigma_z^2$ is set to 1 to match the variance of the standard Gaussian distribution whose probability density function is used to define the probability of motif occurrence in Equation 10 and 11. In the context of our model, $Z_{m,n} > 0$ means that an occurrence of motif $m$ is found in sequence $n$ and is thus equivalent to $A_{m,n} > 0$ while $Z_{m,n} \leq 0$ is equivalent to $A_{m,n} = 0$. This data augmentation scheme is justified by the fact that

$$\begin{aligned}
\pi(Z_{m,n} > 0|y, t, \beta, b, h) &= 1 - \Phi_{\text{mean}=\beta_{m,0}+\sum_c \beta_{m,c}\mathbb{I}\{t_{m,c}=1\}\tilde{y}_{m,n,c},\text{var}=\sigma_z^2}(0) \\
&= 1 - \Phi_{\text{mean}=0,\text{var}=\sigma_z^2}(-\beta_{m,0} - \sum_c \beta_{m,c}\mathbb{I}\{t_{m,c} = 1\}\tilde{y}_{m,n,c}) \\
&= \Phi_{\text{mean}=0,\text{var}=\sigma_z^2}(\beta_{m,0} + \sum_c \beta_{m,c}\mathbb{I}\{t_{m,c} = 1\}\tilde{y}_{m,n,c})\,, \qquad (14)
\end{aligned}$$

where the right-hand side term is the same as in Equation 11 when $\sigma_z^2 = 1$.

#### 1.3 Prior settings and Directed Acyclic Graph of the hierarchical Bayesian model

##### 1.3.1 Priors for the sequence model

As explained in the previous subsection, the starting point of Bayesian inference is the definition of priors for the parameters of the model which will then be considered as random variables. Furthermore, the Bayesian framework allows to treat the variables that tune the dimension of the model as other parameters. To avoid enriching the notations we will use the same notation for the random variable and its value (i.e. not use upper case for the random variable).

The priors that we used intend to be non-informative since we have little *a priori* knowledge about the characteristics of the motifs. Their choice also takes into account considerations on the tractability of the MCMC updates and thus, as usual in Bayesian inference, most of these priors are mutually independent conjugate priors.

The priors for the parameters that affect the dimension of the model are defined as

$$\begin{aligned}
k_m &\sim Geom(\text{probability of success} = p_k)\,, &(15)\\
w_m &\sim Uniform(\{w_{\min}, ..., w_{\max}\})\,, &(16)\\
v &\sim Uniform(\{0, ..., v_{\max}\})\,, &(17)\\
p_m &\sim Bernoulli(p_p)\,, &(18)
\end{aligned}$$

and, when $p_m = 1$,

$$\begin{aligned} \pi(c_m|w_m) &\propto a_c^{|c_m-(w_m+1)/2|} && \text{for } c_m \in \{1.5, 2, 2.5, \ldots, w_m - 0.5\}\,, && (19)\\ q_{m,w}|c_m &\sim \mathit{Bernoulli}(p_q) && \text{for } w \in \{\max(1, 2c_m - w_m), \ldots, \lfloor c_m - 0.5 \rfloor\}\,. && (20) \end{aligned}$$

The emission parameters that describe the nucleotide composition in the background and in the motifs are modelled as drawn from the following independent Dirichlet distributions

$$\begin{aligned} \theta_{0,s} &\sim \mathit{Dirichlet}_4(d_{\theta_0}, d_{\theta_0}, d_{\theta_0}, d_{\theta_0}) \quad \text{for}\, s \in \{\mathtt{A},\mathtt{C},\mathtt{G},\mathtt{T}\}^{v'},\ v' = 0 : v\,, && (21)\\ \theta_{m,w} &\sim \mathit{Dirichlet}_4(d_\theta, d_\theta, d_\theta, d_\theta)\,. && (22) \end{aligned}$$

The priors for the parameters that describe the distribution of the motif occurrences are defined as

$$\begin{aligned} \lambda_m \mid k_m &\sim \mathit{Dirichlet}_{k_m+1}(d_\lambda, \ldots, d_\lambda)\,, && (23)\\ d_m \mid k_m &\sim \mathit{Uniform}(k_m\text{-combinations of } \{2, \ldots, \mathcal{L}\})\,, && (24)\\ r_m \mid w_m &\sim \mathit{Uniform}(\{1, \ldots, w_m\})\,. && (25) \end{aligned}$$

A typical choice that we used in our applications for the values for the parameters of the priors are $w_{\min} = 3$, $w_{\max} = 25$, $v_{\max} = 6$, $p_k = 0.25$, $d_\lambda = 1$, $p_p = 0.5$, $a_c = 0.5$, $p_q = 0.5$, $d_\theta = 0.25$, $d_{\theta_0} = 1$. The purpose of parameter $a_c \leq 1$ is to express a preference for a centre of the palindromic structure near the centre of the PWM. The choice of $p_k$ favours a small $k_m$ and the choice of $d_\theta$ favours PWM columns with high information content.

##### 1.3.2 Priors for the extended probit

We typically have many covariates of type "vector", each summarising one aspect of the expression profiles; but few covariates of type "tree", each providing a global summary of the expression profiles. For this reason, our prior concerning the active covariates (status encoded in variable $t_c$) of the extended probit distinguishes these two types of covariates.

For the covariates of type "tree", we simply use independent Bernoulli priors

$$t_{m,c} \sim \mathit{Bernoulli}(p_{t,\text{tree}})\,. \tag{26}$$

For the covariates of type "vector", we wanted a prior that favours the activation of a small number of covariates which conducted us to use a model in which activation of the different covariates are not independent. Namely, we use a geometric prior on the number of active covariates coupled with a uniform prior on which covariates are active (given the number of active covariates). The corresponding joint density function for the $\mathcal{C}_v$ covariates of type “vector” is

$$\pi((t_{m,c})_{c=1:\mathcal{C}_v}) \quad \propto \quad p_{t,\text{vector}}^{\sum_{c=1:\mathcal{C}_v} t_{m,c}} \frac{1}{\binom{\mathcal{C}_v}{\sum_{c=1:\mathcal{C}_v} t_{m,c}}}\,, \tag{27}$$

in which the sum is over the covariates of type “vector”, and $p_{t,\text{vector}}$ corresponds to the parameter “probability of success” of the geometric prior. From Equation 27, the conditional distribution for any particular $t_{m,c}$ given the status of all other covariates of type “vector” (denoted here $t_{m,-c}$) writes

$$\pi(t_{m,c}|t_{m,-c}) \quad \propto \quad \mathbb{I}\{t_{m,c}=1\}p_{t,\text{vector}}\frac{\mathcal{C}_v - \sum_{-c} t_{m,c'} - 1}{\sum_{-c} t_{m,c'} + 1} + \mathbb{I}\{t_{m,c}=0\}\,, \tag{28}$$

where $\sum_{-c} t_{m,c'}$ corresponds to the number of active components among all covariates of types “vector”, excepted $c$.

The prior used for the parameter $\beta_{m,c}$ are independent Gaussian distributions

$$\beta_{m,0} \quad \sim \quad \mathcal{N}(\text{mean} = m_{\beta_0}, \text{sd} = s_{\beta_0})\,, \tag{29}$$

$$\beta_{m,c} \quad \sim \quad \mathcal{N}(\text{mean} = m_{\beta}, \text{sd} = s_{\beta})\,. \tag{30}$$

A different prior is used for $\beta_{m,0}$ because it correspond to the intercept parameter whose meaning differs from those of the other coefficients of the probit model. In fact, the mean of the prior for the intercept $\beta_{m,0}$ tunes the expected number of occurrences for motif $m$ in absence of link to expression covariates, with 0 corresponding to a presence of the motif in one half of the sequences (see Equation 11). In contrast, $m_\beta$, the mean of the prior for $\beta_{m,c}$, will be set to 0, since the covariates are centred at 0 and have no reason to favour positive or negative links between expression covariates and probability of motif occurrence (the sign of a covariate such as obtained by PCA or ICA is arbitrary).

The prior used for the binarisation of a covariate $c$ of type “vector” is

$$g_{m,c}|t_{m,c} = 1 \quad \sim \quad \mathit{Bernoulli}(p_g)\,. \tag{31}$$

The prior used for cut-off value used in the binarisation process, $h_{m,c}$, which is encoded as rank of the value in the vector or the index of the node in the tree, correspond to a uniform with respect to the values taken by the covariate (in case of covariate of type “vector”) or the length of the tree (in case of covariate of type “tree”). Hence, for a covariate of type “vector”,

$$\pi(h_{m,c}|t_{m,c} = 1, g_{m,c} = 1) \quad \propto \quad y_{[h_{m,c}]+1,c} - y_{[h_{m,c}],c}\,. \tag{32}$$

For a covariate of type “tree”, the prior writes

$$\pi(h_{m,c}|t_{m,c}=1, g_{m,c}=1) \quad \propto \quad (h_{p(h_{m,c}),c} - h_{h_{m,c},c})\mathbb{I}\{n(h_{m,c}) \geq n_{h,\min}, \mathcal{N} - n(h_{m,c}) \geq n_{h,\min}\}\,, \tag{33}$$

where $p(h_{m,c})$ is the parent node of $h_{m,c}$ and $n(h_{m,c})$ is the number of leafs in the subtree hanging below $h_{m,c}$. In this prior, $n_{h,\min}$ is a parameter that avoids locating a breakpoint in the tree at a position that would delineate a singleton or coexpression cluster that we judge too small.

In practice we used $p_{t,\text{tree}} = 0.5$, $p_{t,\text{vector}} = 0.5$, $p_g = 0.5$, $m_{\beta_0} = \Phi^{-1}_{0,1}(0.01)$ (i.e. $m_{\beta_0} \approx -2.326$), $s_{\beta_0} = \sqrt{0.2}$, $m_\beta = 0$, $s_\beta = 1$, and $n_{h,\min} = 10$. The choice of $m_{\beta_0}$ favours motifs with small number of occurrences.

##### 1.3.3 Directed Acyclic Graph

The Directed Acyclic Graph (DAG) represents the factorisation of the joint probability distribution of the variables of our hierarchical model (parameters, hidden variables, observed data). This joint probability distribution writes

$$\begin{aligned}
&\pi(x, a, b, t, \beta, g, h, d, \lambda, k, r, \theta_{1:\mathcal{M}}, q, c, p, w, \theta_0, v|y) \\
&= \pi(x|a, b, r, \theta_{1:\mathcal{M}}, \theta_0)\pi(a|y, t, \beta, g, h, d, \lambda, y)\pi(b|a, r, w, \theta_{1:\mathcal{M}}) \\
&\quad \times\pi(r|w)\pi(d|k)\pi(\lambda|k)\pi(k) \\
&\quad \times\pi(\theta_0|v)\pi(v)\pi(\theta_{1:\mathcal{M}}|q, c, p, w)\pi(q|c, w)\pi(c|p, w)\pi(w) \\
&= \prod_{n,l} \pi(x_{n,l}|x_{n,1:l-1}, a, b, r, \theta_{1:\mathcal{M}}, \theta_0) \prod_{m,n} \pi(a_{m,n}|y, t_m, \beta_m, g_m, h_m, d_m, \lambda_m) \prod_{n,l} \pi(b_{n,l}|a, r, w, \theta_{1:\mathcal{M}}) \\
&\quad \times \prod_m \pi(r_m|w_m) \prod_m \pi(d_m|k_m)\pi(\lambda_m|k_m)\pi(k_m) \\
&\quad \times\pi(\theta_0|v)\pi(v) \prod_m \pi(\theta_m|q_m, c_m, p_m, w_m)\pi(q_m|c_m, w_m)\pi(c_m|p_m, w_m)\pi(w_m)\,, 
\end{aligned} \tag{34}$$

where the term in red is specific to the $\theta$-dependent weight mixture model for overlaps.

The DAG representation helps (via the construction of the corresponding undirected Moral graph in which edges have been added between all pairs of nodes with a common child) to visualise the Markov blanket for each variable and by turn the conditional independence relationships useful for the design of the MCMC steps targeting the different blocks of variables.

The Directed Acyclic Graph presented in Figure 1 encompasses the random variables $t_m$, $\beta_m$, $g_m$, $h_m$, and $Z_m$ that are used to account for expression data.

### 2 MCMC algorithm

#### 2.1 Strategy and overview of the MCMC algorithm

##### 2.1.1 A block MCMC sampler

A Markov Chain Monte Carlo (MCMC) algorithm Robert and Casella (2004); Liu (2004) was built with the joint posterior

$$a, b, v, \theta_0, w, p, c, q, \theta, r, k, d, \lambda, t, \beta, g, h \mid x, y \tag{35}$$

as target distribution. Since the dimension of the target is high, this MCMC algorithm is composed of many different steps whose purposes are to update separate subsets of variables. This type of algorithm is a block MCMC sampler (Andrieu et al., 2003) which consists of a cycle through the different steps such as to allow the update of all the variables. Each step is designed to preserve the target distribution by sampling from the conditional distribution of a block of variables given all the other variables. Usually, the conditional independence properties make that conditional distributions involve only a subset of the other variables. In some cases, variables also disappear from the sampled conditional distribution because they are marginalised out. A subsequent step is then needed to "restore" the status of these variables that were marginalised out by sampling from their conditional distribution.

The steps of the block MCMC sampler that we implemented can be conveniently divided into three types:

- Metropolis-Hastings steps (abbreviated MH steps) that rely on a proposal coupled to an accept/reject procedure to preserve a target distribution (i.e. generating a Markov chain with the target as stationary distribution if repeated);
- Gibbs steps that consists of sampling directly from the target (conditional distribution of the variable or block of variables);
- Reversible-Jump MH steps that are generalisation of the Metropolis-Hastings steps when the attempted moves change the dimension (Green, 1995).

Gibbs step can be seen as a special case of Metropolis-Hastings step where the proposal coincides with the target conditional distribution. Similarly, under circumstances where the probability distribution of the variables whose dimension is modified can be integrated-out, the Reversible-Jump MH steps can be done in a Gibbs manner, i.e. direct sampling from the conditional distribution of the variable that defines the dimension. We refer to this particular case as dimension-changing Gibbs step.

##### 2.1.2 Overview of the 14 steps of the MCMC algorithm

In total, 14 steps have been designed taking into account the structure of dependence summarised in the DAG (Figure 1) and computational considerations. Using the notation $\theta$ for $(\theta_0, (\theta_m)_{m=1:\mathcal{M}})$ and in their order of appearance in the implemented sweeps, these steps that compose our algorithm are:

1. Update the palindromic structure of motif $m$, described by $(p_m, c_m, q_m)$ according to $p_m, c_m, q_m \mid a_m, b, r_m, w_m, x$ (Dimension-changing Gibbs step, $\theta_m$ is marginalised out). Details are provided page 34.

2. Update the PWM $\theta_m$, either according to $\theta_m \mid a, b, r, w, p_m, c_m, q_m, x$ (Gibbs step) if the equal weight mixture model for overlaps is used; or preserving $\theta_m \mid \theta_{-m}, a, b, r, w, p_m, c_m, q_m, x$ (MH step) if the $\theta$-dependent weight mixture model for overlaps is used. Here, we use the notation $\theta_{-m}$ to refer to all the components of $\theta = (\theta_0, \theta_1, \dots, \theta_{\mathcal{M}})$ but $\theta_m$ (the same notation will also be used for other variables). Note that this update restores the status of $\theta_m$ which was marginalised out when updating the palindromic structure. Details are provided page 24.

3. Update the data augmentation variable $z_{m,n}$ of the probit model according to $z_{m,n} \mid a_{m,n}, t_m, \beta_m, g_m, h_m$ (Gibbs step). Details are provided page 36.

4. Update the dimension changing variables $t_{m,c}, g_{m,c}, h_{m,c}$ according to $t_{m,c}, g_{m,c}, h_{m,c} \mid z_m, \beta_{m,-c}, t_{m-c}, g_{m,-c}, h_{m,-c}$ (dimension changing Gibbs step in which $\beta_{m,c}$ is marginalised out). Details are provided page 38.

5. Update the coefficients of the probit regression $\beta_{m,c}$ according to $\beta_{m,c} \mid z_m, \beta_{m,-c}, t_m, g_m, h_m$ (Gibbs step). Details are provided page 37.

6. Update the position of the occurrences of the motif $m$, $a_m$, according to $a_m \mid a_{-m}, t_m, \beta_m, g_m, h_m, \theta, r, y, t, \beta, g, h, x$ (Gibbs step, $b$ and $z_m$ are marginalised out). Details are provided page 23.

7. Update motif width $w_m$, preserving $w_m, \theta_m, r_m, c_m, q_m \mid a, r_{-m}, \theta_{-m}, p_m, x$ (Reversible-Jump MH step, $b$ is marginalised out). Details are provided page 27.

8. Update the PWM $\theta_m$ by shifting motif $m$ , preserving $\theta_m, r_m, c_m, q_m \mid a, r_{-m}, \theta_{-m}, p_m, x$ (Reversible-Jump MH step, with joint update of $\theta_m$ and $r_m$, $b$ is marginalised out). This update which consists of removing one column of the PWM on a side and adding a column on the other side, is simply the coupling of two updates of motif width (decrease on left side coupled with increase on right side,

decrease on right side coupled with increase on left side) whose details are provided page 27. It was introduced to allow the motif occurrences to "slide" along the sequences even when $w_m = w_{\max}$.

9. Update the disambiguation variables for motif overlaps, $b$, according to $b \mid a, r, \theta, x$ (Gibbs step). Note that this update restores the status of $b$ which was marginalised out when updating $a$ and $w$. Details are provided page 24.

10. Update the Markov order of the background, $v$, preserving $v \mid a, r, w, x$ (Reversible-Jump step, $\theta_0$ is marginalised out). Details are provided page 30.

11. Update the nucleotide composition of the background, $\theta_0$, according to $\theta_0 \mid v, a, r, w, x$ (Gibbs step). Note that this update restores the status of $\theta_0$ which was marginalised out when updating $v$. Details are provided page 29.

12. Update the position of the breakpoints defining the piece-wise constant probability density function modelling the positions of occurrences of motif $m$, $d_m$, preserving $d_m \mid k_m, a_m$ (MH step, $\lambda$ is marginalised out). Details are provided page 32.

13. Update the number of breakpoints in the piece-wise constant probability density function, $k_m$, preserving $k_m, d_m \mid a_m$ (Reversible-Jump MH step, note the joint update of $d$, while $\lambda$ is marginalised out). Details are provided page 33.

14. Update the expected fraction of motif occurrences found in each region of the piece-wise constant pdf, $\lambda$, according to $\lambda_m \mid k_m, d_m, a_m$ (Gibbs step). Note that this step restores the status of $\lambda$ which was marginalised out when updating $d$ and $k$. Details are provided page 32.

15. Update the reference positions motif $m$, $r_m$, preserving $r_m, a_m \mid w_m, d_m, \lambda_m$ (MH step, note the joint update of $a$). Details are provided page 31.

Of note, this ordering of the steps makes that the variables that were marginalised out are restored before their reuse. Other orders would work provided that marginalised-out variables are appropriately restored (which may induce extra-computation).

The 3 steps that update the variables needed for the implementation of the extended probit model ($z_{m,n}$, $\beta_{m,c}$, $h_{m,c}$, $t_{m,c}$, $g_{m,c}$) are relatively fast (in particular compared to the update of $a$) but they consider each covariate $c$ separately and they are done conditionally on $a_{m,n}$ which may slow the mixing. This sequence of 3 steps is thus repeated 10 times at each sweep of the algorithm in our MCMC runs.

##### 2.1.3 Detecting errors in the MCMC algorithm

Design and implementation of an MCMC algorithms with so many steps is an error prone process.

In order to detect errors in the equations or implementation of our MCMC algorithm we also implemented a step consisting in simulating sequence $x$ given all the other variables (i.e. $x \mid a, b, \theta$). When activated the MCMC algorithm targets the full joint distribution

$$x, a, b, t, \beta, g, h, d, \lambda, k, r, \theta_{1:\mathcal{M}}, q, c, p, w, \theta_0, v \mid y$$

instead of the conditional

$$a, b, t, \beta, g, h, d, \lambda, k, r, \theta_{1:\mathcal{M}}, q, c, p, w, \theta_0, v \mid x, y\,.$$

The marginal distribution of the parameters under this full joint distribution corresponds to the injected priors and many any errors in the implementation or in the equations skew these marginals and can thus be detected by comparing the marginals to the priors (Geweke, 2004). For this purpose we typically work in a space of small dimension (e.g. taking $\mathcal{N} = 10$ and $\mathcal{M} = 2$) which considerably speeds up the convergence and thereby allows very precise comparisons of the marginals to the priors after a relatively short run (typical length between 1 and 30 minutes).

#### 2.2 Details of 14 the steps of MCMC algorithm

Instead of providing the details of the steps in their order of appearance in one sweep of the MCMC algorithm, we adopt here an order that intends to make the presentation easier to understand. We start by describing the update of the hidden variables $a$ and $b$ and then continue by describing, for each group of variables related to a same aspect of the model, the simple updates before the dimension-changing updates.

##### 2.2.1 Update of $a$, the position of the occurrences of the motifs

This update is conducted as a Gibbs step, separately for each motif and each sequence, in which $b$ is marginalised out. The conditional density of $a_{m,n}|a_{-m,n}, \theta, r, y, t, \beta, g, h, x$, the position of motif $m$ in sequence $n$, is computed, up to a normalising constant, at each point of the support $\{0, 1, \ldots, \mathcal{L}\}$ using the formula

$$\begin{aligned}
\pi(a_{m,n}|a_{-m,n},\theta,y,t,\beta,g,h,r,x) &= \frac{\pi(a_{m,n},a_{-m,n},\theta,y,t,\beta,g,h,r,x)}{\pi(a_{-m,n},\theta,y,t,\beta,g,h,r,x)} \\
&\propto \pi(x_n|a_{m,n},a_{-m,n},\theta,r)\pi(a_{m,n}|y,t,\beta,g,h) \\
&\propto \left[\prod_{l=a_{m,n}-r_m+1}^{a_{m,n}-r_m+w_m} \frac{\pi(x_{n,l}|a_{m,n},a_{-m,n},\theta,r)}{\pi(x_{n,l}|A_{m,n}=0,a_{-m,n},\theta,r)}\right]^{\mathbb{I}\{a_{m,n}\neq 0\}} \\
&\quad \times\pi(a_{m,n}|y,t,\beta,g,h)\,,
\end{aligned} \tag{36}$$

where the term $\pi(a_{m,n}|y,t,\beta,g,h)$ is given by Equation 9 and the term of the form $\pi(x_{n,l}|a_{m,n} = k, a_{-m,n}, \theta, r)$ are obtained by summing over all possible values of $b_{n,l}$

$$\begin{aligned}
\pi(x_{n,l}|a,\theta,r) &= \sum_{m=1:\mathcal{M}} \pi(x_{n,l}|B_{n,l}=m,a,\theta,r)\pi(B_{n,l}=m|a,\theta,r)\,, \\
&= \sum_{m=1:\mathcal{M}} \theta_{m,l-(a_{m,n}-r_m)+1,x_{n,l}}\pi(B_{n,l}=m|a,\theta,r)\,,
\end{aligned} \tag{37}$$

where $\pi(b_{n,l}|a, \theta, r)$ is given by Equation 7 or 8 (depending of the type of mixture model chosen for overlaps). The new value of $a_{m,n}$ is then drawn directly according to the conditional density computed at the $\mathcal{L} + 1$ points of the support.

##### 2.2.2 Update of $b$, the disambiguation variables for motif overlaps

This update is done successively for all positions $(n, l)$ by direct sampling from the conditional density of $b_{n,l}|a, r, \theta, x$ (Gibbs step) obtained as

$$\pi(b_{n,l}|a,r,\theta,x) \propto \pi(x_{n,l}|b_{n,l},a,r,\theta)\pi(b_{n,l}|a,r,\theta)\,,$$

where the two types of terms are given by Equations 4 and by Equations 7 or 8, depending of the type of mixture model chosen for motif overlaps. The new value of $b_{n,l}$ is then drawn directly according to the conditional density computed at the $\mathcal{M} + 1$ points of the support. Of note, this conditional distribution is trivial if zero or only one motif covers the position $(n, l)$, with probability of 1 for $b_{n,l} = 0$ if zero motif, and probability of 1 for $b_{n,l} = m$ if only motif $m$.

##### 2.2.3 Update of $\theta_m$, the PWM of motif $m$

As already stated, this update can consists of a Gibbs or a MH step depending on the type of mixture model chosen for motif overlaps (equal weight vs. $\theta$-dependent weight). In all cases, the parameters of the different $m$ and of the different columns $w$ within each motif are updated successively (excepted for columns paired in a palindromic structure that requires simultaneous update) and the conditional distribution $\theta_m|\theta_{-m}, a, b, r, w, p_m, c_m, q_m, x$ is preserved.

Let's first describe the simple Gibbs step for the equal weight mixture. In this case, we need to distinguish the case of a column $(m, w)$ that is not paired in a palindromic structure and the case of a paired column. The first case arises when the motif is not palindromic ($p_m = 0$), or the column cannot be paired due to its position relative to the centre of the palindrome ($w < 2c_m - w_m$, $w = c_m$ or $w \geq 2c_m$), or the column could have been paired but is not paired ($q_{m,min(w,2c_m-w)} = 0$). In this first case, the conditional distribution writes

$$\begin{aligned} &\theta_{m,w}|a, b, r, w, p_m, c_m, q_{m,min(w,2c_m-w)} = 0, x \\ &\sim \; Dirichlet_4(\ldots, d_\theta + c_{m,w,u}, \ldots)\,, \end{aligned} \tag{38}$$

where $c_{m,w,u}$ is the count of the nucleotide $u$ emitted from column $(m, w)$ defined as

$$\begin{aligned} c_{m,w,u} &= \sum_{n,l} \mathbb{I}\{x_{n,l} = u, b_{n,l} = m, a_{m,n} - r_m + w = l\} \\ &= \sum_{n} \mathbb{I}\{x_{n,a_{m,n}-r_m+w} = u, b_{n,l} = m\}\,. \end{aligned} \tag{39}$$

If the column is paired ($p_m = 1$ and $q_{m,\min(w,2c_m-w)} = 1$), then $\theta_{m,w,u} = \theta_{m,2c_m-w,\bar{u}}$. This implies the simultaneous update of the two PWM columns ($w$ and $2c_m - w$) by a single drawing from the conditional distribution obtained by aggregating the counts from the two paired columns

$$\begin{aligned} &\theta_{m,w} \mid a, b, r, w, p_m, c_m, q_{m,w} = 1, x \\ &\sim \; Dirichlet_4(\ldots, d_\theta + c_{m,w,u} + c_{m,2c_m-w,\bar{u}}, \ldots)\,. \end{aligned} \tag{40}$$

Equation 38 is a direct and well known (Neuwald et al., 1995) consequence of the choice of a conjugate prior for $\theta_{m,w}$. From Equations 4 and 22 and using Bayes' rule, this conditional distribution is obtained as follows

$$\begin{aligned} \pi(\theta_{m,w}|x, \ldots) &\propto \pi(\theta_{m,w}, x, \ldots) \\ &\propto \pi(x|\theta_{m,w}, \ldots)\pi(\theta_{m,w}|\ldots) \\ &\propto \prod_{n,l}\prod_{u} \theta_{m,w,u}^{\mathbb{I}\{x_{n,l}=u, b_{n,l}=m, a_{m,n}-r+w=l\}} \times \prod_{u} \theta_{m,w,u}^{d_\theta - 1} \\ &\propto \prod_{u} \theta_{m,w,u}^{d_\theta + \sum_{n,l}\mathbb{I}\{x_{n,l}=u, b_{n,l}=m, a_{m,n}-r+w=l\} - 1}\,, \end{aligned} \tag{41}$$

where "…" is a convenient notation to refer to all the other variables (i.e. here besides $x$ and $\theta_{m,w}$). Using the notation $c_{m,w,u}$ defined in Equation 39, the normalised version of this conditional density writes

$$\pi(\theta_{m,w}|x,\ldots) = \frac{\Gamma(\sum_u d_\theta + c_{w,m,u})}{\prod_u \Gamma(d_\theta + c_{w,m,u})} \prod_u \theta_{m,w,u}^{d_\theta + c_{w,m,u} - 1}, \tag{42}$$

and corresponds to the Dirichlet distribution of Equation 38. The same line of reasoning leads to Equation 40 for paired columns in palindromic motifs.

The update of $\theta_m$ when the overlap model is a $\theta$-dependent mixture is more complicated due to the grey arrow linking from $\theta_m$ to $b$ in the DAG (Figure 1). The analogous of Equation 41 which gives the conditional density of $\theta_{m,w}$ is then

$$\begin{aligned}\pi(\theta_{m,w}|x,b,\ldots) &\propto \pi(\theta_{m,w},x,b,\ldots)\\ &\propto \pi(x|\theta_{m,w},b,\ldots)\pi(b|\theta_{m,w},\ldots)\pi(\theta_{m,w}|\ldots)\\ &\propto \pi(b|\theta_{m,w},\ldots)\prod_u \theta_{m,w,u}^{d_\theta + \sum_{n,l}\mathbb{I}\{x_{n,l}=u, b_{n,l}=m, a_{m,n}-r+w=l\}-1},\end{aligned} \tag{43}$$

where "..." refer to all the variables besides $x$, $b$ and $\theta_{m,w}$. This conditional density can no longer be identified to the density of a Dirichlet distribution which precludes direct sampling (Gibbs step).

The workaround that we implemented is to build a MH step using the Dirichlet of Equation 38 as a proposal. In simple words, we propose a new value according to the conditional distribution under the simple model and use an accept-reject method to match it to the conditional under the more complex model. Denoting $q_{b,a,x}$ the density of this proposal, the probability of acceptation of the proposed value $\tilde{\theta}_{m,n}$ writes

$$\alpha(\theta_{m,n},\tilde{\theta}_{m,n}) = \min\left\{1; \frac{\pi(\tilde{\theta}_{m,w}|b,\ldots)q_{b,a,x}(\theta_{m,w})}{\pi(\theta_{m,w}|b,\ldots)q_{b,a,x}(\tilde{\theta}_{m,w,u})}\right\}. \tag{44}$$

Using Equation 43, this probability of acceptation simplifies to

$$\begin{aligned}\alpha(\theta_{m,n},\tilde{\theta}_{m,n}) &= \min\left\{1; \frac{\pi(b|\tilde{\theta}_{m,w},\ldots)}{\pi(b|\theta_{m,w},\ldots)}\right\}\\ &= \min\left\{1; \prod_{n,l}\left(\frac{\pi(b_{n,l}|\tilde{\theta}_{m,w},\ldots)}{\pi(b_{n,l}|\theta_{m,w},\ldots)}\right)^{\mathbb{I}\{a_{m,n}\neq 0, l=a_{m,n}-r_m+w\}}\right\},\end{aligned} \tag{45}$$

which involves to compute a product whose number of terms equals to the number of occurrences of motif $m$ where column $w$ is overlapped by an occurrence of another motif (the terms corresponding to the other occurrences are equal to 1). Each of these individual terms is easy to obtain from Equation 8.

##### 2.2.4 Update of $w_m$, the width of motif $m$

This update relies on a Reversible-Jump MH step in which a new values $\tilde{w}_m, \tilde{\theta}_m, \tilde{r}_m, \tilde{c}_m, \tilde{q}_m$ are proposed for $w_m, \theta_m, r_m, c_m, q_m$. Importantly, the update does not change the position of the motifs ($a$ is given) and each motif are treated successively. The variable $b$ is marginalised out. The update preserves the conditional distribution $w_m, \theta_m, r_m, c_m, q_m | a, r_{-m}, \theta_{-m}, p_m, x$.

The proposed move consists of adding or removing only one column, i.e. $\tilde{w}_m = w_m + 1$ or $\tilde{w}_m = w_m - 1$. Four "directions" of modification of the PWM are possible depending on the sign (increase or decrease the width of the PWM) and on the side (modification on the right side or on the left side of the PWM). In order to preserve the position of the occurrences and since $a$ is kept unchanged, if the modification is done on the left side the reference position needs to be shifted of one bp, i.e. $\tilde{r}_m = r_m + \tilde{w}_m - w_m$. Of note, this implies that a decrease on the left side is automatically rejected if $r_m = 1$ and a decrease on the right side is automatically rejected if $r_m = w_m$. If the modification is done on the left side, $r_m$ is unchanged.

The update of the width preserves palindromic structures ($p_m$ is given). To preserve the pairing of the columns, this implies, if the change is done on the left side of a palindromic motif, to shift the centre of the palindrome $c_m$ and the pairing variables $q_{m,w}$, i.e. $\tilde{c}_m = c_m + \tilde{w}_m - w_m$ and $\tilde{q}_{m,w+\tilde{w}_m - w_m} = q_{m,w}$ for $2c_m - w_m \leq w < c_m$ provided that $w + \tilde{w}_m - w_m \geq 0$. This design of proposal means that we cannot accept a modification of the width that would shift the centre of the palindrome outside of its allowed range, which happens when $c_m \in \{1.5, 2\}$ and a decrease of the width is attempted on the left side or $c_m \in \{w_m - 1, w_m - 0.5\}$ and a decrease of the width is attempted on the right side. To cope with the palindromic structure, we also need to define a proposal for the pairing status of the added column when an increase of the motif width is proposed and the palindromic structure defined by $(p_m, w_m, c_m)$ allows the pairing of this new column, which happens with left increase when $2c_m < w_m + 1$ and right increase $2c_m > w_m + 1$ (this pairing concerns $\tilde{q}_{m,1}$ if left increase and $\tilde{q}_{2\tilde{c}_m - \tilde{w}_m}$ if right increase). For simplicity, we decided that the new column will be unpaired (i.e. $\tilde{q}_{m,1} = 0$ or $\tilde{q}_{2\tilde{c}_m - \tilde{w}_m} = 0$). A direct consequence of this choice is to forbid decrease moves that remove paired columns (this constraint can be seen in the ratio of proposal in the acceptation probability, Equation 49).

Our proposal gives the same probability 1/4 to each of the four directions of modification (left vs. right, increase vs. decrease) and it keeps unchanged the values of $\theta_{m,w}$ for those columns that are not directly concerned by the change. Namely, for a change on the right side, we have thus $\tilde{\theta}_m = (\theta_m, \tilde{\theta}_{m,w_m+1})$ if $\tilde{w}_m = w_m + 1$ and $\tilde{\theta}_m = (\theta_{m,-w_m})$ if $\tilde{w}_m = w_m - 1$. Similarly, for a change on the left side, $\tilde{\theta}_m = (\tilde{\theta}_{m,1}, \theta_m)$ if $\tilde{w}_m = w_m + 1$ and $\tilde{\theta}_m = (\theta_{m,-1})$ if $\tilde{w}_m = w_m - 1$. The proposal for a move that increases the width involves thus the drawing of a single four dimensional vector $\tilde{\theta}_{m,w}$ corresponding to the added PWM column. To maximise the chance of accepting the proposed modification, our proposal takes into account the nucleotide composition of the positions in the sequence that will be overlapped by this new PWM column if the move is accepted as follow

$$\begin{aligned}\tilde{\theta}_{m,w_m+1} &\sim Dirichlet_4(\ldots,\max\{d_\theta+\tilde{c}_{m,w_m+1,r}-\tilde{o}_{m,w_m+1,r},1\},\ldots) \quad \text{if increase right}\\ \tilde{\theta}_{m,1} &\sim Dirichlet_4(\ldots,\max\{d_\theta+\tilde{c}_{m,-1,r}-\tilde{o}_{m,-1,r},1\},\ldots) \quad \text{if increase left,}\end{aligned} \tag{46}$$

where $\tilde{c}_{m,w,u}$ corresponds to the count of nucleotide $r$ at the sequence positions covered by the new column $w$ (differing from $c_{m,w,u}$ of Equation 39 by the fact that $b$ is not taken into account), and $\tilde{o}_{m,w,u}$ corresponds to the expected contribution of the other motifs that overlap the occurrences of motif $m$ to the count of nucleotide $r$ (according to the equal weight mixture model for motif overlap). Equation 46 sets the lower boundary of the parameters of the Dirichlet to 1. This avoids (very rare) cases were $d_\theta + \tilde{c}_{m,w,u} - \tilde{o}_{m,w,u}$ is negative and is therefore incompatible with the range of values allowed for the parameters of a Dirichlet. The count $\tilde{c}_{m,w,u}$ is obtained as

$$\tilde{c}_{m,w,u} = \sum_n \mathbb{I}\{a_{m,n}\neq 0, x_{n,a_{m,n}-r_m+w}=r\}\,. \tag{47}$$

and the expected count $\tilde{o}_{m,w,u}$ as

$$\begin{aligned}\tilde{o}_{m,w,u} = \sum_{n,m'\neq m} &\Bigg\{\mathbb{I}\{a_{m,n}\neq 0, x_{n,a_{m,n}-r_m+w}=r\}\theta_{m',a_{m,n}-r_m+w-a_{m',n}-r_{m'},r}\\ &\times\frac{\mathbb{I}\{a_{m',n}\neq 0, 1\leq a_{m,n}-r_m+w-a_{m',n}-r_{m'}\leq w_{m'}\}}{\sum_{m''}\mathbb{I}\{a_{m'',n}\neq 0, 1\leq a_{m,n}-r_m+w-a_{m'',n}-r_{m''}\leq w_{m''}\}}\Bigg\}.\end{aligned} \tag{48}$$

Denoting by $q$ the proposal described in the preceding paragraphs, the acceptation ratio of the Reversible-Jump MH move writes

$$\begin{aligned}&\alpha((w_m,\theta_m,r_m,c_m,q_m),(\tilde{w}_m,\tilde{\theta}_m,\tilde{r}_m,\tilde{c}_m,\tilde{q}_m))\\ &= \min\left\{1;\frac{\pi(\tilde{w}_m,\tilde{\theta}_m,\tilde{r}_m,\tilde{c}_m,\tilde{q}_m|a,r_{-m},\theta_{-m},x)q(w_m,\theta_m,r_m,c_m,q_m)}{\pi(w_m,\theta_m,r_m,c_m,q_m|a,r_{-m},\theta_{-m},x)q(\tilde{w}_m,\tilde{\theta}_m,\tilde{r}_m,\tilde{c}_m,\tilde{q}_m)}\right\}\\ &= \min\left\{1;\frac{\pi(x|\tilde{w}_m,\tilde{\theta}_m,\tilde{r}_m,a,r_{-m},\theta_{-m})}{\pi(x|w_m,\theta_m,r_m,a,r_{-m},\theta_{-m})}\times\frac{\pi(\tilde{w}_m,\tilde{\theta}_m,\tilde{r}_m,\tilde{c}_m,\tilde{q}_m)}{\pi(w_m,\theta_m,r_m,c_m,q_m)}\right.\\ &\qquad\left.\times\frac{q(w_m,\theta_m,r_m,c_m,q_m)}{q(\tilde{w}_m,\tilde{\theta}_m,\tilde{r}_m,\tilde{c}_m,\tilde{q}_m)}\right\}.\end{aligned} \tag{49}$$

Of note, many terms simplify in this ratio. To illustrate its computation, we can take the example of an attempt to increase the width of the motif on the right side (i.e. $\tilde{w}_m = w_m + 1$,

$\tilde{r}_m = r_m$, and $\tilde{\theta}_m = (\theta_m, \tilde{\theta}_{m,w_m+1})$). In this case, the probability of acceptation

$$\alpha((w_m, \theta_m, r_m, c_m, q_m), (\tilde{w}_m, \tilde{\theta}_m, \tilde{r}_m, \tilde{c}_m, \tilde{q}_m)) \quad = \quad \min\{1, A\}\,, \tag{50}$$

is build on acceptation ratio

$$\begin{aligned} A \quad = \quad & \prod_{n:a_{m,n}\neq 0} \frac{\pi(x_{n,a_{m,n}-r_m+w_m+1}|\tilde{w}_m, \tilde{\theta}_m, \tilde{r}_m, a, r_{-m}, \theta_{-m})}{\pi(x_{n,a_{m,n}-r_m+w_m+1}|w_m, \theta_m, r_m, a, r_{-m}, \theta_{-m})} \times \frac{\pi(\tilde{c}_m, \tilde{q}_m|p_m)}{\pi(c_m, q_m|p_m)} \\ & \times \frac{w_m}{w_m+1} \times \frac{Dir_4(\tilde{\theta}_{m,w_m+1}; \ldots, d_\theta, \ldots)}{Dir_4(\tilde{\theta}_{m,w_m+1}; \ldots, \max\{d_\theta + \tilde{c}_{m,-1,r} - \tilde{o}_{m,-1,r}, 1\}, \ldots)}\,, \end{aligned} \tag{51}$$

where the term $w_m/(w_m+1)$ corresponds to $\pi(\tilde{r}_m|\tilde{w}_m)/\pi(r_m|w_m)$. In this formula, the terms of the form $\pi(x_{n,l}|w_m, \theta_m, r_m, a, r_{-m}, \theta_{-m})$ are obtained by summing over all possible values of $b_{n,l}$ (see Equation 37). The reverse modification from $(\tilde{w}, \tilde{\theta})$ to $(w, \theta)$ that decreases the width on the right side has probability of acceptation $\min\{1, 1/A\}$.

##### 2.2.5 Update of $\theta_0$, the nucleotide composition of the background

The choice of a product of independent 4-dimensional Dirichlet priors for the parameter $\theta_0$ of the Markov model of order $v$ describing the background composition of the sequences (outside motifs) makes that the posterior is also a product of independent 4-dimensional Dirichlet distributions (property of conjugate prior). This allows direct sampling from the conditional distribution $\theta_0 \mid v, a, r, w, x$ (Gibbs step).

When we include the initial distribution (i.e. transitions for orders $0 \leq k < v$ that serve to model the first positions of the sequences), the number of Dirichlet distributions is $4^0 + 4^1 + \ldots + 4^v = (4^{v+1} - 1)/3$, since $\theta_0 = (((\theta_{0,s,u})_{u\in\{\mathtt{A},\mathtt{C},\mathtt{G},\mathtt{T}\}})_{s\in\{\mathtt{A},\mathtt{C},\mathtt{G},\mathtt{T}\}^{v'}})_{v'=0:v}$.

Using the notation $o_{n,l}$ for the number of motif occurrences that overlap position $(n, l)$ defined in Equation 6, the conditional density of $\theta_0$ decomposes as the following product of conditional densities for each $\theta_{0,s}$,

$$\begin{aligned} \pi(\theta_0|v, a, r, w, x) \quad & \propto \quad \pi(x|\theta_0, a, r, w)\pi(\theta_0|v) \\ & \propto \quad \pi(\theta_0|v) \prod_{n,l} \pi(x_{n,l}|x_{n,max(1,l-v):l-1}, \theta_0, a, r, w)^{\mathbb{I}\{o_{n,l}=0\}} \\ & \propto \quad \prod_{s\in(\{\mathtt{A},\mathtt{C},\mathtt{G},\mathtt{T}\}^{v'})_{v'=0:v}} \underbrace{\pi(\theta_{0,s}) \prod_{u\in\{\mathtt{A},\mathtt{C},\mathtt{G},\mathtt{T}\}} \theta_{0,s,u}^{\sum_{n,l} \mathbb{I}\{o_{n,l}=0\, x_{n,max(1,l-v):l-1}=s, x_{n,l}=u\}}}_{\propto\, \pi(\theta_{0,s}|v,a,r,w,x)}\,. \end{aligned} \tag{52}$$

The conditional density $\pi(\theta_{0,s}|v,a,r,w,x)$ can be rewritten, up to a normalising constant, as

$$\pi(\theta_{0,s}|v,a,r,w,x) \quad \propto \quad \prod_{u\in\{\mathtt{A},\mathtt{C},\mathtt{G},\mathtt{T}\}} \theta_{0,s,u}^{d_{\theta_0}+\sum_{n,l}\mathbb{I}\{o_{n,l}=0\, x_{n,max(1,l-v-1):l-1}=s,x_{n,l}=u\}-1}, \quad (53)$$

which can be identified to the density of the Dirichlet distribution that we use to update $\theta_{0,s}$,

$$\begin{aligned}&\theta_{0,s}|v,a,r,w,x \\ &\quad\sim \quad \mathit{Dirichlet}_4(\ldots, d_{\theta_0} + \sum_{n,l} \mathbb{I}\{o_{n,l}=0\, x_{n,max(1,l-v):l-1}=s, x_{n,l}=r\}, \ldots)\,.\end{aligned} \quad (54)$$

##### 2.2.6 Update of $v$, the Markov order of the background

This dimension changing update is carried out as Reversible-Jump MH step preserving $v \mid a,r,w,x$ in which $\theta_0$ is marginalised out.

Given the current Markov order $v$, a new value $\tilde{v} \in \{v-1, v+1\}$ is proposed, according to proposal $q_v$ (in practice we use $q_v(v+1)=0.5$ and $q_v(v-1)=0.5$). This new value is accepted with probability

$$\begin{aligned}\alpha(v,\tilde{v}) &= \min\left\{1; \frac{\pi(\tilde{v}|a,r,w,x)q_{\tilde{v}}(v)}{\pi(v|a,r,w,x)q_v(\tilde{v})}\right\} \\ &= \min\left\{1; \frac{\pi(x|\tilde{v},a,r,w)}{\pi(x|v,a,r,w)} \times \frac{\pi(\tilde{v})}{\pi(v)} \times \frac{q_{\tilde{v}}(v)}{q_v(\tilde{v})}\right\} \\ &= \min\left\{1; \frac{\prod_{n,l}\pi(x_{n,l}|x_{n,max(1,l-\tilde{v}):l-1},\tilde{v},a,r,w)^{\mathbb{I}\{o_{n,l}=1\}}}{\prod_{n,l}\pi(x_{n,l}|x_{n,max(1,l-v):l-1},v,a,r,w)^{\mathbb{I}\{o_{n,l}=1\}}} \times \frac{\pi(\tilde{v})}{\pi(v)} \times \frac{q_{\tilde{v}}(v)}{q_v(\tilde{v})}\right\},\end{aligned} \quad (55)$$

where $o_{n,l}$ corresponds to the number of motifs overlapping position $(n,l)$ as defined in Equation 6. The products of conditional densities in which $\theta_0$ is marginalised out that appear in the probability of acceptation can be obtained in close form. Using the notation $c_{0,s,r}^{(v)} = \sum_{n,l}\mathbb{I}\{o_{n,l}=0, x_{n,max(1,l-v-1):l-1}=s, x_{n,l}=r\}$ for the count of word $(s,r)$ with last character $r$ in the background, these products are computed as

$$\begin{aligned}
&\prod_{n,l} \pi(x_{n,l}|x_{n,max(1,l-v):l-1}, v, a, r, w)^{\mathbb{I}\{o_{n,l}=1\}} \\
&= \int_{\theta_0} \pi(\theta_0) \prod_{n,l} \pi(x_{n,l}|x_{n,max(1,l-v):l-1}, \theta_0, v, a, r, w)^{\mathbb{I}\{o_{n,l}=1\}} d\theta_0 \\
&= \prod_{s\in(\{\mathtt{A,C,G,T}\}^{v'})_{v'=0:v}} \int_{\theta_{0,s}} \pi(\theta_{0,s}) \prod_{u\in\{\mathtt{A,C,G,T}\}} \theta_{0,s,u}^{c^{(v)}_{0,s,u}} d\theta_{0,s} \\
&= \prod_{s\in(\{\mathtt{A,C,G,T}\}^{v'})_{v'=0:v}} \frac{\Gamma(\sum_{u\in\{\mathtt{A,C,G,T}\}} d_{\theta_0})}{\prod_{u\in\{\mathtt{A,C,G,T}\}} \Gamma(d_{\theta_0})} \int_{\theta_{0,s}} \prod_{u\in\{\mathtt{A,C,G,T}\}} \theta_{0,s,u}^{d_{\theta_0}+c^{(v)}_{0,s,u}-1} d\theta_{0,s} \\
&= \left(\frac{\Gamma(\sum_{u\in\{\mathtt{A,C,G,T}\}} d_{\theta_0})}{\prod_{u\in\{\mathtt{A,C,G,T}\}} \Gamma(d_{\theta_0})}\right)^{(4^{v+1}-1)/3} \prod_{s\in(\{\mathtt{A,C,G,T}\}^{v'})_{v'=0:v}} \frac{\Gamma(\sum_{u\in\{\mathtt{A,C,G,T}\}} d_{\theta_0} + c^{(v)}_{0,s,u})}{\prod_{u\in\{\mathtt{A,C,G,T}\}} \Gamma(d_{\theta_0} + c^{(v)}_{0,s,u})} .
\end{aligned} \tag{56}$$

##### 2.2.7 Update of $r_m$, the reference positions motif $m$.

This update consists of a simple MH step preserving $r_m, a_m \mid w_m, d_m, \lambda_m$ in which new values $\tilde{r}_m$ and $\tilde{a}_m$ are proposed for $r_m$ and $a_m$. In practice, an attempt is made to increase or decrease $r_m$ by 1 (with equal probabilities) and, simultaneously, to shift the positions of the motif encoded in $a_m$ such as to maintain the positions of the occurrences of the motifs. Namely,

$$\tilde{a}_{m,n} \quad = \quad a_{m,n} + (\tilde{r}_m - r_m)\mathbb{I}\{a_{m,n} \neq 0\} \quad \text{for } n = 1 : \mathcal{N} . \tag{57}$$

Importantly, when shifting the $a_{m,n}$, we do not allow motif occurrences to disappear, i.e. are moved outside of the range $\{1, \dots, \mathcal{L}\}$. Therefore, the attempted move is automatically rejected when $\tilde{r}_m = r_m - 1$ if there exists $(m, n)$ such as $a_{m,n} = 1$ and $\tilde{r}_m = r_m + 1$ if there exists $(m, n)$ such as $a_{m,n} = \mathcal{L}$.

The probability of acceptation for this attempted move writes as

$$\begin{aligned}
\alpha((r_m, a_m), (\tilde{r}_m, \tilde{a}_m)) &= \min\left\{1; \frac{\pi(\tilde{r}_m, \tilde{a}_m|x, \dots)q_{\tilde{r}_m,\tilde{a}_m}(r_m, a_m)}{\pi(r_m, a_m|x, \dots)q_{r_m,a_m}(\tilde{r}_m, \tilde{a}_m)}\right\} \\
&= \min\left\{1; \frac{\pi(x|\tilde{r}_m, \tilde{a}_m, \dots)\pi(\tilde{r}_m| \dots)\pi(\tilde{a}_m \mid \dots)q_{\tilde{r}_m,\tilde{a}_m}(r_m, a_m)}{\pi(x|r_m, a_m, \dots)\pi(r_m| \dots)\pi(a_m \mid \dots)q_{r_m,a_m}(\tilde{r}_m, \tilde{a}_m)}\right\} \\
&= \min\left\{1; \frac{\pi(\tilde{r}_m|w_m)\pi(\tilde{a}_m \mid d_m, \lambda_m)}{\pi(r_m|w_m)\pi(a_m \mid d_m, \lambda_m)}\right\} ,
\end{aligned} \tag{58}$$

where the ratio $\pi(\tilde{r}_m|w_m)/\pi(r_m|w_m)$ is 1 if $1 \leq \tilde{r}_m \leq w_m$ and 0 otherwise, and the terms of the form $\pi(a_m \mid d_m, \lambda_m)$ are given by Equation 9.

##### 2.2.8 Update of $\lambda_m$, the expected fraction of motif occurrences found in each region of the piecewise constant pdf

The expected fraction of motif occurrences found in each region of the piecewise constant pdf, $\lambda_m = (\lambda_{m,1}, \ldots, \lambda_{m,k_m+1})$ are updated for each motif $m$ by direct drawing from the conditional density $\lambda_m \mid k_m, d_m, a_m$. Such a Gibbs step is allowed by the choice of Dirichlet distribution as prior for $\lambda_m | k_m$ (property of conjugate prior). From Equation 9,

$$\begin{aligned}
\pi(\lambda_m | k_m, d_m, a_m) &\propto \pi(\lambda_m | k_m) \pi(a_m | d_m, k_m) \\
&\propto \prod_{k=1:k_m+1} \lambda_{m,k}^{d_\lambda - 1} \prod_n \lambda_{m,k}^{\mathbb{I}\{d_{m,k-1} \leq a_{m,n} < d_{m,k}\}} \\
&\propto \prod_{k=1:k_m+1} \lambda_{m,k}^{d_\lambda + \sum_n \mathbb{I}\{d_{m,k-1} \leq a_{m,n} < d_{m,k}\} - 1}, \qquad (59)
\end{aligned}$$

which corresponds to the density of the Dirichlet

$$\lambda_m | k_m, d_m, a_m \sim Dirichlet_{k_m+1}(\ldots, d_\lambda + c_{m,k}, \ldots), \qquad (60)$$

using the notation

$$c_{m,k} = \sum_n \mathbb{I}\{d_{m,k-1} \leq a_{m,n} < d_{m,k}\}. \qquad (61)$$

##### 2.2.9 Update of $d_m$, the positions of the breakpoints defining the piecewise constant pdf modelling the positions of occurrences of motif $m$

The positions $d_m = (d_{m,1}, \ldots, d_{m,k})$ of the $k_m$ breakpoints defining the piecewise constant pdf modelling the positions of occurrences of motif $m$ are updated by a MH step preserving $d_m \mid k_m, a_m$ in which $\lambda_m$ is marginalised out. The proposed $\tilde{d}_m$ is obtained by choosing one of the $k_m$ breakpoints and assigning it a new position in $\{2, \ldots, \mathcal{L}\}$ among the $\mathcal{L} - k_m$ positions not occupied by the $k_m - 1$ unchanged breakpoints. This proposal corresponds to an uniform distribution over $k_m(\mathcal{L} - k_m)$ distinct $\tilde{d}_m$ that can be obtained by changing the position of one of the breakpoints of $d_m$,

$$q_{d_m}(\tilde{d}_m) = \frac{1}{k_m(\mathcal{L} - k_m)}. \qquad (62)$$

The MH acceptation ratio simplifies due to the symmetry of the proposal ($q_{d_m}(\tilde{d}_m) = q_{\tilde{d}_m}(d_m)$) and the uniform prior on $d_m$ over all the $k_m$-combinations in $\{2, \dots, \mathcal{L}\}$,

$$\begin{aligned}
\alpha(d_m, \tilde{d}_m) &= \min\left\{1; \frac{\pi(\tilde{d}_m|k_m, a_m, y, t_m, \beta_m, g_m, h_m) q_{\tilde{d}_m}(d_m)}{\pi(d_m|k_m, a_m, y, t_m, \beta_m, g_m, h_m) q_{d_m}(\tilde{d}_m)}\right\} \\
&= \min\left\{1; \frac{\pi(\tilde{d}_m|k_m)\pi(a_m|\tilde{d}_m, y, t_m, \beta_m, g_m, h_m) q_{\tilde{d}_m}(d_m)}{\pi(d_m|k_m)\pi(a_m|d_m, y, t_m, \beta_m, g_m, h_m) q_{d_m}(\tilde{d}_m)}\right\} \\
&= \min\left\{1; \frac{\pi(a_m|\tilde{d}_m, y, t_m, \beta_m, g_m, h_m)}{\pi(a_m|d_m, y, t_m, \beta_m, g_m, h_m)}\right\} \qquad (63)
\end{aligned}$$

where the terms of the form $\pi(a_m|d_m, y, t_m, \beta_m, g_m, h_m)$ are obtained by marginalising out $\lambda_m$, which is made possible by the Dirichlet prior on this parameter. From Equation 9 and using the notation $c_{m,k}$ defined in Equation 61, we obtain

$$\begin{aligned}
&\pi(a_m|d_m, y, t_m, \beta_m, g_m, h_m) \\
&= \int_{\lambda_m} \pi(a_m|\lambda_m, d_m, y, t_m, \beta_m, g_m, h_m)\pi(\lambda_m|d_m) d\lambda_m \\
&= \frac{\prod_n \alpha_{m,n}^{\mathbb{I}\{a_{m,n}\neq 0\}}(1-\alpha_{m,n})^{\mathbb{I}\{a_{m,n}=0\}}}{\prod_{k=1:k_m+1}(d_{m,k}-d_{m,k-1})^{c_{m,k}}} \times \frac{\prod_{k=1:k_m+1}\Gamma(d_\lambda)}{\Gamma(\sum_{k=1:k_m+1} d_\lambda)} \times \int_{\lambda_m} \prod_{k=1:k_m+1} \lambda_{m,k}^{d_\lambda + c_{m,k}-1} d\lambda_m \\
&\propto \frac{1}{\prod_{k=1:k_m+1}(d_{m,k}-d_{m,k-1})^{c_{m,k}}} \times \frac{\prod_{k=1:k_m+1}\Gamma(d_\lambda)}{\Gamma(\sum_{k=1:k_m+1} d_\lambda)} \times \frac{\Gamma(\sum_{k=1:k_m+1} d_\lambda + c_{m,k})}{\prod_{k=1:k_m+1}\Gamma(d_\lambda + c_{m,k})}. \qquad (64)
\end{aligned}$$

Using this closed form for the marginalised density, the acceptation ratio of Equation 63 writes

$$\begin{aligned}
\alpha(d_m, \tilde{d}_m) = \min\Bigg\{1; &\frac{\prod_{k=1:k_m+1}(d_{m,k}-d_{m,k-1})^{c_{m,k}}}{\prod_{k=1:k_m+1}(\tilde{d}_{m,k}-\tilde{d}_{m,k-1})^{\tilde{c}_{m,k}}} \\
&\times \frac{\Gamma(\sum_{k=1:k_m+1} d_\lambda + \tilde{c}_{m,k}) \prod_{k=1:k_m+1}\Gamma(d_\lambda + c_{m,k})}{\Gamma(\sum_{k=1:k_m+1} d_\lambda + c_{m,k}) \prod_{k=1:k_m+1}\Gamma(d_\lambda + \tilde{c}_{m,k})}\Bigg\}, \qquad (65)
\end{aligned}$$

where $\tilde{c}_{m,k}$ is the analogous of $c_{m,k}$ defined using $\tilde{d}_m$ instead of $d_m$.

###### 2.2.10 Update of $k_m$, the number of breakpoints in the piecewise constant pdf

The update of $k_m$ is a Reversible-Jump MH step based on the proposition of a couple $(\tilde{k}_m, \tilde{d}_m)$ whose acceptation probability preserves the conditional distribution $k_m, d_m|a_m$ ($\lambda_m$ is marginalised out). This update is very similar to the update of $d_m$ (page 32) but consists of adding a removing (instead of moving) a breakpoint in $d_m$. When the addition of a new breakpoint is proposed (i.e. $\tilde{k}_m = k_m + 1$) its position is selected randomly among the $\mathcal{L} - k_m - 1$ positions in $\{2, \dots, \mathcal{L}\}$ that are not occupied by the $k_m$ already existing breakpoints. The proposal for the move that decreases $k_m$ consists of removing a randomly selecting and removing one breakpoint. The proposal density associated to $(\tilde{k}_m, \tilde{d}_m)$ generated by this procedure writes thus

$$q_{k_m,d_m}(\tilde{k}_m, \tilde{d}_m) = \frac{1}{2k_m}\mathbb{I}\{\tilde{k}_m = k_m - 1\} + \frac{1}{2(\mathcal{L} - k_m - 1)}\mathbb{I}\{\tilde{k}_m = k_m + 1\} . \tag{66}$$

Analogously to Equation 63, the probability of acceptation decomposes into

$$\alpha((k_m, d_m), (\tilde{k}_m, \tilde{d}_m)) = \min\left\{1; \frac{\pi(\tilde{k}_m)\pi(\tilde{d}_m|\tilde{k}_m)\pi(a_m|\tilde{d}_m, y, t_m, \beta_m, g_m, h_m)q_{\tilde{k}_m,\tilde{d}_m}(k_m, d_m)}{\pi(k_m)\pi(d_m|k_m)\pi(a_m|d_m, y, t_m, \beta_m, g_m, h_m)q_{k_m,d_m}(\tilde{k}_m, \tilde{d}_m)}\right\} . \tag{67}$$

Taking the example of an increase move (i.e. $\tilde{k}_m = k_m+1$) and using the result of Equation 64, this probability can be written

$$\begin{aligned}
&\alpha((k_m, d_m), (\tilde{k}_m, \tilde{d}_m)) \\
&= \min\left\{1; (1 - p_k)\frac{\binom{\mathcal{L}-1}{k_m}}{\binom{\mathcal{L}-1}{k_m+1}} \times \frac{\pi(a_m|\tilde{d}_m, y, t_m, \beta_m, g_m, h_m)}{\pi(a_m|d_m, y, t_m, \beta_m, g_m, h_m)} \times \frac{\mathcal{L} - k_m - 1}{k_m}\right\} \qquad (68) \\
&= \min\left\{1; (1 - p_k)\frac{k_m + 1}{k_m}\right. \\
&\qquad \times \frac{\prod_{k=1:k_m+1}(d_{m,k} - d_{m,k-1})^{c_{m,k}}}{\prod_{k=1:\tilde{k}_m+1}(\tilde{d}_{m,k} - \tilde{d}_{m,k-1})^{\tilde{c}_{m,k}}} \times \frac{\Gamma(d_\lambda)\Gamma(\sum_{k=1:k_m+1} d_\lambda)}{\Gamma(\sum_{k=1:\tilde{k}_m+1} d_\lambda)} \\
&\qquad \left. \times \frac{\Gamma(\sum_{k=1:\tilde{k}_m+1} d_\lambda + \tilde{c}_{m,k}) \prod_{k=1:k_m+1} \Gamma(d_\lambda + c_{m,k})}{\Gamma(\sum_{k=1:k_m+1} d_\lambda + c_{m,k}) \prod_{k=1:\tilde{k}_m+1} \Gamma(d_\lambda + \tilde{c}_{m,k})}\right\} ,
\end{aligned} \tag{69}$$

where $\tilde{c}_{m,k}$ and $c_{m,k}$ are defined by Equation 61.

###### 2.2.11 Update of $(p_m, c_m, q_m)$, the palindromic structure of motif $m$

The use of Dirichlet priors for the columns of $\theta_m$ allows, in the case of the equal weight mixture model for overlaps, to sample directly from the conditional distribution $p_m, c_m, q_m|a_m, b, r_m, w_m, x$ ($\theta_m$ is marginalised out). Of note, this Gibbs-type update changes the dimension of the model. Such a joint update is very efficient but could not be implemented for the $\theta$-dependent mixture model for overlaps which precludes the use of this model when palindromic structures are allowed (alternative updates have not been implemented). In practice, sampling from $p_m, c_m, q_m|a, b, r_m, w_m, x$ is done in two steps:

- sample from $p_m, c_m|a_m, b, r_m, w_m, x$ (marginalising out $q_m$),

- sample from $q_m|c_m, p_m, a_m, b, r_m, w, x$.

The densities needed for these two steps are obtained by appropriate summing and normalisation (for marginalisation and conditioning) of the joint conditional density

$$\begin{aligned}
&\pi(p_m, c_m, q_m|a, b, r, w_m, x) \\
&\quad \propto \quad \pi(p_m, c_m, q_m|w_m)\pi(x|a, b, r, w_m, p_m, c_m, q_m) \\
&\quad \propto \quad \pi(p_m, c_m, q_m|w_m) \\
&\qquad\qquad \times \underbrace{\pi\big((x_{n,a_{m,n}-r_m+w})_{(n,w):a_{m,n}\neq 0, w\in\{1,\ldots,w_m\}, b_{n,a_{m,n}-r_m+w}=m}|a, b, r, w_m, p_m, c_m, q_m\big)}_{L(p_m,c_m,q_m)},
\end{aligned} \tag{70}$$

where $L(p_m, c_m, q_m)$ is a likelihood term that decomposes in a product of $w_m$ terms. If $p_m = 0$ (no palindromic structure), $c_m$ and $q_m$ are not defined and the likelihood can be written

$$\begin{aligned}
L(p_m = 0, c_m = \emptyset, q_m = \emptyset) &= \prod_{w=1:w_m} \int_{\theta_{m,w}} \theta_{m,w}^{c_{m,w,u}} \pi(\theta_{m,w}) d\theta_{m,w} \\
&= \prod_{w=1:w_m} \frac{\Gamma(\sum_u d_\theta)}{\prod_u \Gamma(d_\theta)} \int_{\theta_{m,w}} \theta_{m,w}^{c_{m,w,u}+d_\theta-1} d\theta_{m,w} \\
&= \prod_{w=1:w_m} \frac{\Gamma(\sum_u d_\theta)}{\prod_u \Gamma(d_\theta)} \frac{\prod_u \Gamma(d_\theta + c_{m,w,u})}{\Gamma(\sum_u d_\theta + c_{m,w,u})}.
\end{aligned} \tag{71}$$

When $p_m = 1$ (palindromic structure), we need to group the paired columns (where $\theta_{m,w,u} = \theta_{m,2c_m-w,\bar{u}}$) to marginalise over $\theta_{m,w}$, which gives

$$\begin{aligned}
&L(p_m = 0, c_m, q_m) \\
&\quad = \prod_{w=1:w_m} \left\{ \left[ \frac{\Gamma(\sum_u d_\theta)}{\prod_u \Gamma(d_\theta)} \frac{\prod_u \Gamma(d_\theta + c_{m,w,u})}{\Gamma(\sum_u d_\theta + c_{m,w,u})} \right]^{\mathbb{I}\{w<2c_m-w_m\}+\mathbb{I}\{w=c_m\}+\mathbb{I}\{w\geq 2c_m\}} \right. \\
&\qquad \times \left[ \mathbb{I}\{q_{m,w} = 0\} \left( \frac{\Gamma(\sum_u d_\theta)}{\prod_u \Gamma(d_\theta)} \right)^2 \frac{\prod_u \Gamma(d_\theta + c_{m,w,u})}{\Gamma(\sum_u d_\theta + c_{m,w,u})} \frac{\prod_u \Gamma(d_\theta + c_{m,2c_m-w,r})}{\Gamma(\sum_u d_\theta + c_{m,2c_m-w,r})} \right. \\
&\qquad\qquad \left. \left. + \mathbb{I}\{q_{m,w} = 1\} \frac{\Gamma(\sum_u d_\theta)}{\prod_u \Gamma(d_\theta)} \frac{\prod_u \Gamma(d_\theta + c_{m,w,u} + c_{m,2c_m-w,\bar{u}})}{\Gamma(\sum_u d_\theta + c_{m,w,u} + c_{m,2c_m-w,\bar{u}})} \right]^{\mathbb{I}\{2c_m-w_m\leq w<c_m\}} \right\}.
\end{aligned} \tag{72}$$

Using the formulas of Equations 71 and 72 for the likelihood terms, we obtain the density needed for the step 1 sampling $(p_m, c_m)$ by summing the joint density of Equation 70 over all possible values of $q_m$,

$$\begin{aligned}
&\pi(p_m, c_m|a, b, r, w, x) \\
&\quad\propto \sum_{q_m} \pi(q_m, c_m, p_m|a, b, r, w, x) \\
&\quad\propto \pi(p_m)\pi(c_m|p_m) \sum_{q_m} \pi(q_m|c_m) L(p_m, c_m, q_m) \\
&\quad\propto \pi(p_m)\pi(c_m|p_m) \prod_{w=1:w_m} \left\{ \left[ \frac{\Gamma(\sum_u d_\theta)}{\prod_u \Gamma(d_\theta)} \frac{\prod_u \Gamma(d_\theta + c_{m,w,u})}{\Gamma(\sum_u d_\theta + c_{m,w,u})} \right]^{\mathbb{I}\{w<2c_m-w_m\}+\mathbb{I}\{w=c_m\}+\mathbb{I}\{w\geq 2c_m\}} \right. \\
&\qquad\qquad \times \left[ (1-p_q) \left( \frac{\Gamma(\sum_u d_\theta)}{\prod_u \Gamma(d_\theta)} \right)^2 \frac{\prod_u \Gamma(d_\theta + c_{m,w,u})}{\Gamma(\sum_u d_\theta + c_{m,w,u})} \frac{\prod_u \Gamma(d_\theta + c_{m,2c_m-w,r})}{\Gamma(\sum_u d_\theta + c_{m,2c_m-w,r})} \right. \\
&\qquad\qquad\quad \left. \left. + p_q \frac{\Gamma(\sum_u d_\theta)}{\prod_u \Gamma(d_\theta)} \frac{\prod_u \Gamma(d_\theta + c_{m,w,u} + c_{m,2c_m-w,\bar{u}})}{\Gamma(\sum_u d_\theta + c_{m,w,u} + c_{m,2c_m-w,\bar{u}})} \right]^{\mathbb{I}\{2c_m-w_m\leq w<c_m\}} \right\}. \qquad (73)
\end{aligned}$$

For those columns that can be paired (i.e. $w$ such as $\min(1, 2c_m - w_m) \leq w < c_m$), the conditional density used in the step 2 of the update (sampling $q_{m,w}$ given $c_m$) is obtained from Equations 70, 72, and 20 as

$$\begin{aligned}
&\pi(q_{m,w}|c_m, p_m, a, b, r, w, x) \\
&\quad\propto \mathbb{I}\{q_{m,w}=0\}(1-p_q) \left( \frac{\Gamma(\sum_u d_\theta)}{\prod_u \Gamma(d_\theta)} \right)^2 \frac{\prod_u \Gamma(d_\theta + c_{m,w,u})}{\Gamma(\sum_u d_\theta + c_{m,w,u})} \frac{\prod_u \Gamma(d_\theta + c_{m,2c_m-w,r})}{\Gamma(\sum_u d_\theta + c_{m,2c_m-w,r})} \\
&\qquad + \mathbb{I}\{q_{m,w}=1\} p_q \frac{\Gamma(\sum_u d_\theta)}{\prod_u \Gamma(d_\theta)} \frac{\prod_u \Gamma(d_\theta + c_{m,w,u} + c_{m,2c_m-w,\bar{u}})}{\Gamma(\sum_u d_\theta + c_{m,w,u} + c_{m,2c_m-w,\bar{u}})}.
\end{aligned} \qquad (74)$$

##### 2.2.12 Update of $z$, the data augmentation variable of probit model

The $z_{m,n}$'s are updated successively by direct sampling from the conditional distribution of $z_{m,n}$ given $a_{m,n}, \beta_m, t_m, g_m, h_m$ (Gibbs step). This distribution is a truncated Gaussian since

$$\begin{aligned}
&\pi(z_{m,n}|a_{m,n}, \beta_m, t_m, g_m, h_m, \ldots) \\
&\quad\propto \pi(a_{m,n}|z_{m,n}, \beta_m, t_m, g_m, h_m, \ldots)\pi(z_{m,n}|\beta_m, t_m, g_m, h_m) \\
&\quad\propto \left( \mathbb{I}\{a_{m,n} > 0, z_{m,n} \geq 0\} + \mathbb{I}\{a_{m,n} = 0, z_{m,n} < 0\} \right) \\
&\qquad \times \exp\left\{ -\frac{1}{2\sigma_z^2} \left( z_{m,n} - \beta_0 - \sum_c \mathbb{I}\{t_{m,c}=1\}\beta_{m,c}\tilde{y}_{m,n,c} \right)^2 \right\}. \qquad (75)
\end{aligned}$$

Hence, given that $\sigma_z^2 = 1$ (see Equation 13, but is still written as $\sigma_z^2$ to make apparent the homogeneity of the formulas), we draw $z_{m,n}$ from $\mathcal{N}(\text{mean} = \beta_0 + \sum_c \mathbb{I}\{t_{m,c}=1\}\beta_{m,c}\tilde{y}_{m,n,c}, \text{var} = 1)$ truncated to $\mathbb{R}^-$ if $a_{m,n} = 0$ and truncated to $\mathbb{R}^+$ if $a_{m,n} > 0$.

##### 2.2.13 Update of $\beta$, the coefficients of probit regression

The $\beta_{m,c}$ are updated successively for the active covariates ($t_{m,c} = 1$) by direct sampling from the conditional distribution of $\beta_{m,c}$ given $z_m, \beta_{m,-c}, t_m, g_m, h_m$. This update is made possible by the choice of a Gaussian prior for $\beta$ which is the conjugate prior for the mean of a Gaussian distribution. The possibility of such a Gibbs step is the fundamental purpose of the introduction of $z$. Of note, we update here each $\beta_{m,c}$ separately for simplicity but a joint update would also be possible.

The conditional distribution of $\beta_{m,c}$ is easy to write up to a constant,

$$\begin{aligned}
&\pi(\beta_{m,c}|z_m, \beta_{m,-c}, t_m, g_m, h_m) \\
&\quad \propto \ \pi(z_m|\beta_{m,c}\beta_{m,-c}, t_m, g_m, h_m)\pi(\beta_{m,c}) \\
&\quad \propto \ \prod_n \exp\Big\{-\frac{1}{2\sigma_z^2}\big(z_{m,n} - \beta_0 - \sum_{c'} \mathbb{I}\{t_{m,c'} = 1\}\beta_{m,c'}\tilde{y}_{m,n,c'}\big)^2\Big\} \\
&\qquad\quad \times \exp\Big\{-\frac{1}{2s_\beta^2}(\beta_{m,c} - m_\beta)^2\Big\} \\
&\quad \propto \ \exp\Bigg\{-\frac{1}{2s_\beta^2}(\beta_{m,c} - m_\beta)^2 \\
&\qquad\quad -\frac{1}{2\sigma_z^2}\sum_n \Big(z_{m,n} - \beta_0 - \mathbb{I}\{t_{m,c} = 1\}\beta_{m,c}\tilde{y}_{m,n,c} - \sum_{c' \neq c} \mathbb{I}\{t_{m,c'} = 1\}\beta_{m,c'}\tilde{y}_{m,n,c'}\Big)^2\Bigg\} \\
&\quad \propto \ \exp\Bigg\{-\frac{1}{2}\beta_{m,c}^2\Bigg[\frac{1}{s_\beta^2} + \frac{\mathbb{I}\{t_{m,c} = 1\}}{\sigma_z^2}\sum_n \tilde{y}_{n,c}^2\Bigg] \\
&\qquad\quad -\beta_{m,c}\Bigg[\frac{m_\beta}{s_\beta^2} + \frac{\mathbb{I}\{t_{m,c} = 1\}}{\sigma_z^2}\sum_n \tilde{y}_{m,n,c}\Big(z_{m,n} - \beta_0 - \sum_{c' \neq c} \mathbb{I}\{t_{m,c'} = 1\}\beta_{m,c'}\tilde{y}_{m,n,c'}\Big)\Bigg]\Bigg\} .
\end{aligned} \tag{76}$$

For the conciseness of the subsequent formulas, we will use the notation

$$z_{m,n,-c} \ = \ z_{m,n} - \beta_0 - \sum_{c' \neq c} \mathbb{I}\{t_{m,c'} = 1\}\beta_{m,c'}\tilde{y}_{m,n,c'} \, . \tag{77}$$

The right-hand side term of Equation 76 corresponds to the density of the Gaussian distribution that we use to update $\beta_{m,c}$,

$$\pi(\beta_{m,c}|z_m, \beta_{m,-c}, t_m, g_m, h_m) \ \sim \ \mathcal{N}(\text{mean} = \mu_{\beta_{m,c}|\ldots}, \text{var} = \sigma^2_{\beta_{m,c}|\ldots}) \, , \tag{78}$$

where

$$\mu_{\beta_{m,c}|\ldots} = \left[\frac{m_\beta}{s_\beta^2} + \frac{\mathbb{I}\{t_{m,c}=1\}}{\sigma_z^2}\sum_n \tilde{y}_{m,n,c} z_{m,n,-c}\right] \times \left[\frac{1}{s_\beta^2} + \frac{\mathbb{I}\{t_{m,c}=1\}}{\sigma_z^2}\sum_n \tilde{y}_{n,c}^2\right]^{-1}$$

$$\sigma^2_{\beta_{m,c}|\ldots} = \left[\frac{1}{s_\beta^2} + \frac{\mathbb{I}\{t_{m,c}=1\}}{\sigma_z^2}\sum_n \tilde{y}_{n,c}^2\right]^{-1}. \quad (79)$$

The update of $\beta_{m,0}$ relies on analogous equations in which the $\tilde{y}_{m,n,c}$'s are replaced by 1,

$$\pi(\beta_{m,0}|z_m, \beta_{m,-0}, t_m, g_m, h_m) \sim \mathcal{N}(\text{mean} = \mu_{\beta_{m,0}|\ldots}, \text{var} = \sigma^2_{\beta_{m,0}|\ldots}), \quad (80)$$

with

$$\mu_{\beta_{m,0}|\ldots} = \left[\frac{m_{\beta_0}}{s_{\beta_0}^2} + \frac{1}{\sigma_z^2}\sum_n \left(z_{m,n} - \sum_c \mathbb{I}\{t_{m,c}=1\}\beta_{m,c}\tilde{y}_{m,n,c}\right)\right] \times \left[\frac{1}{s_{\beta_0}^2} + \frac{\mathcal{N}}{\sigma_z^2}\right]^{-1}$$

$$\sigma^2_{\beta_{m,0}|\ldots} = \left[\frac{1}{s_{\beta,0}^2} + \frac{\mathcal{N}}{\sigma_z^2}\right]^{-1}. \quad (81)$$

##### 2.2.14 Update of the dimension changing variables $t_{m,c}, g_{m,c}, h_{m,c}$ of the extended probit

The possibility to carry out at a relatively small computational cost this update in a Gibbs manner (i.e. by direct sampling of the conditional) is a key ingredient of the usefulness of the extended probit model. This part of the algorithm consists of the joint update of the three variables $t_{m,c}, g_{m,c}, h_{m,c}$ successively for each motif $m$ and selected number of covariates $c$. In practice, we randomly select one tenth of the covariates of type "vector" and one covariate of type "tree". The variable $\beta_{m,c}$ is marginalised out.

The conditional density needed for this update decomposes in three terms corresponding to the mutually exclusive cases of a covariate $c$ which is not active ($t_{m,c} = 0$), a covariate $c$ which is active and not binarised ($t_{m,c} = 1, g_{m,c} = 0$), and a covariate $c$ which is active and binarised ($t_{m,c}=1, g_{m,c}=1$). Namely, we can write

$$
\begin{aligned}
&\pi(t_{m,c}, g_{m,c}, h_{m,c}|z_m, \beta_{m,-c}, t_{m,-c}, g_{m,-c}, h_{m,-c})\\
&\quad\propto\ \pi(z_m|t_{m,c}, g_{m,c}, h_{m,c}, \beta_{m,-c}, t_{m,-c}, g_{m,-c}, h_{m,-c})\pi(t_{m,c}, g_{m,c}, h_{m,c}|t_{m,-c})\\
&\quad\propto\ \mathbb{I}\{t_{m,c}=0\}\pi(t_{m,c}|t_{m,-c})\pi(z_m|t_m, \beta_{m,-c}, g_{m,-c}, h_{m,-c}, t_{m,c}=0)\\
&\qquad+\mathbb{I}\{t_{m,c}=1, g_{m,c}=0\}\pi(t_{m,c}|t_{m,-c})\pi(g_{m,c}|t_{m,c})\\
&\qquad\quad\times\int_{\beta_{m,c}}\pi(z_m|t_m, g_m, \beta_m, h_{m,-c}, t_{m,c}=1, g_{m,c}=0)\pi(\beta_{m,c}|t_{m,c})d\beta_{m,c}\\
&\qquad+\mathbb{I}\{t_{m,c}=1, g_{m,c}=1\}\pi(t_{m,c}|t_{m,-c})\pi(g_{m,c}, h_{m,c}|t_{m,c})\\
&\qquad\quad\times\int_{\beta_{m,c}}\pi(z_m|t_{m,c}, g_{m,c}, h_m, \beta_m, t_{m,c}=1, g_{m,c}=1)\pi(\beta_{m,c}|t_{m,c})d\beta_{m,c}\,.
\end{aligned}
\tag{82}
$$

We will now see how these three terms can be computed, and in particular the third term which needs to be computed efficiently for every possible value of $h_{m,c}$ to be able to draw directly from the conditional distribution of $t_{m,c}, g_{m,c}, h_{m,c}$. The term needed for the first case ($t_{m,c}=0$) writes simply

$$
\begin{aligned}
\pi(z_m|t_m, \beta_{m,-c}, g_{m,-c}, h_{m,-c}, t_{m,c}=0) &= \prod_n \frac{1}{\sigma_z\sqrt{2\pi}}\exp\Big\{-\frac{1}{2\sigma_z^2}z_{m,n,-c}^2\Big\}\\
&= \frac{1}{\sigma_z^{\mathcal{N}}(2\pi)^{\mathcal{N}/2}}\exp\Big\{-\frac{\sum_n z_{m,n,-c}^2}{2\sigma_z^2}\Big\},
\end{aligned}
\tag{83}
$$

where $z_{m,n,-c}$ corresponds to the definition given by Equation 77.

The term needed for the second case ($t_{m,c}=1, g_{m,c}=0$) is obtained as follow

$$\begin{aligned}
&\pi(z_m|t_m,\beta_{m,-c},g_{m,-c},h_{m,-c},t_{m,c}=1,g_{m,c}=0)\\
&= \int_{\beta_{m,c}} \pi(z_m|t_m,g_m,\beta_m,h_{m,-c},t_{m,c}=1,g_{m,c}=0)\pi(\beta_{m,c}|t_{m,c})d\beta_{m,c}\\
&= \int_{\beta_{m,c}} \Big[\prod_n \frac{1}{\sigma_z\sqrt{2\pi}}\exp\Big\{-\frac{1}{2\sigma_z^2}\big(z_{m,n,-c}-\beta_{m,c}y_{n,c}\big)^2\Big\}\\
&\qquad\qquad \times\frac{1}{s_\beta\sqrt{2\pi}}\exp\Big\{-\frac{1}{2s_\beta^2}(\beta_{m,c}-m_\beta)^2\Big\}\Big]d\beta_{m,c}\\
&= \frac{1}{s_\beta\sigma_z^{\mathcal{N}}(2\pi)^{(\mathcal{N}+1)/2}}\times\exp\Big\{-\frac{\sum_n z_{m,n,-c}^2}{2\sigma_z^2}-\frac{m_\beta^2}{2s_\beta^2}\Big\}\\
&\quad\times\int_{\beta_{m,c}}\exp\Big\{-\frac{1}{2}\beta_{m,c}^2\Big(\frac{\sum_n y_{n,c}^2}{\sigma_z^2}+\frac{1}{s_\beta^2}\Big)+\beta_{m,c}\Big(\frac{\sum_n y_{n,c}z_{m,n,-c}}{\sigma_z^2}+\frac{m_\beta}{s_\beta^2}\Big)\Big\}d\beta_{m,c}\\
&= \frac{1}{s_\beta\sigma_z^{\mathcal{N}}(2\pi)^{\mathcal{N}/2}}\exp\Big\{-\frac{\sum_n z_{m,n,-c}^2}{2\sigma_z^2}\Big\}\times\Big(\frac{\sum_n y_{n,c}^2}{\sigma_z^2}+\frac{1}{s_\beta^2}\Big)^{-1/2}\\
&\quad\times\exp\Big\{-\frac{m_\beta^2}{2s_\beta^2}+\frac{1}{2}\Big(\frac{\sum_n y_{n,c}z_{m,n,-c}}{\sigma_z^2}+\frac{m_\beta}{s_\beta^2}\Big)^2\Big(\frac{\sum_n y_{n,c}^2}{\sigma_z^2}+\frac{1}{s_\beta^2}\Big)^{-1}\Big\}.
\end{aligned}\tag{84}$$

The derivation of the term needed for the third case ($t_{m,c}=1, g_{m,c}=1$) is similar to Equation 84, excepted that $\tilde{y}_{m,n,c}$ replaces $y_{n,c}$. However, it is important to note that $\tilde{y}_{m,n,c}$ depends on $h_{m,c}$ (see Equation 12) and will therefore be written here $\tilde{y}_{m,n,c,h_{m,c}}$ to make this dependence apparent.

$$\begin{aligned}
&\pi(z_m|t_m,\beta_{m,-c},g_{m,-c},h_m,t_{m,c}=1,g_{m,c}=1)\\
&= \frac{1}{s_\beta\sigma_z^{\mathcal{N}}(2\pi)^{\mathcal{N}/2}}\exp\Big\{-\frac{\sum_n z_{m,n,-c}^2}{2\sigma_z^2}\Big\}\times\Big(\frac{\sum_n \tilde{y}_{n,c,h_{m,c}}^2}{\sigma_z^2}+\frac{1}{s_\beta^2}\Big)^{-1/2}\\
&\quad\times\exp\Big\{-\frac{m_\beta^2}{2s_\beta^2}+\frac{1}{2}\Big(\frac{\sum_n \tilde{y}_{m,n,c,h_{m,c}}z_{m,n,-c}}{\sigma_z^2}+\frac{m_\beta}{s_\beta^2}\Big)^2\Big(\frac{\sum_n \tilde{y}_{n,c,h_{m,c}}^2}{\sigma_z^2}+\frac{1}{s_\beta^2}\Big)^{-1}\Big\}.
\end{aligned}\tag{85}$$

Using the expressions obtained for the three terms in Equations 83, 84, and 85, we can rewrite the conditional joint density of $t_{m,c}, g_{m,c}, h_{m,c}$ (Equation 82) as

$$
\begin{aligned}
&\pi(t_{m,c}, g_{m,c}, h_{m,c}|z_m, \beta_{m,-c}, t_{m,-c}, g_{m,-c}, h_{m,-c}) \\
&\quad\propto\quad \pi(z_m|t_{m,c}, g_{m,c}, h_{m,c}, \beta_{m,-c}, t_{m,-c}, g_{m,-c}, h_{m,-c})\pi(t_{m,c}, g_{m,c}, h_{m,c}|t_{m,-c}) \\
&\quad\propto\quad \mathbb{I}\{t_{m,c}=0\}\pi(t_{m,c}|t_{m,-c}) \\
&\qquad\quad +\mathbb{I}\{t_{m,c}=1, g_{m,c}=0\}\pi(t_{m,c}|t_{m,-c})\pi(g_{m,c}|t_{m,c}) \\
&\qquad\quad \Big(\frac{\sum_n y_{n,c}^2}{\sigma_z^2}+\frac{1}{s_\beta^2}\Big)^{-1/2} \\
&\qquad\qquad \times\exp\Big\{-\frac{m_\beta^2}{2s_\beta^2}+\frac{1}{2}\Big(\frac{\sum_n y_{n,c}z_{m,n,-c}}{\sigma_z^2}+\frac{m_\beta}{s_\beta^2}\Big)^2\Big(\frac{\sum_n y_{n,c}^2}{\sigma_z^2}+\frac{1}{s_\beta^2}\Big)^{-1}\Big\} \\
&\qquad\quad +\mathbb{I}\{t_{m,c}=1, g_{m,c}=1\}\pi(t_{m,c}|t_{m,-c})\pi(g_{m,c}|t_{m,c})\pi(h_{m,c}|t_{m,c}, g_{m,c}) \\
&\qquad\qquad \times\Big(\frac{\sum_n \tilde{y}_{n,c,h_{m,c}}^2}{\sigma_z^2}+\frac{1}{s_\beta^2}\Big)^{-1/2} \\
&\qquad\qquad \times\exp\Big\{-\frac{m_\beta^2}{2s_\beta^2}+\frac{1}{2}\Big(\frac{\sum_n \tilde{y}_{m,n,c,h_{m,c}}z_{m,n,-c}}{\sigma_z^2}+\frac{m_\beta}{s_\beta^2}\Big)^2\Big(\frac{\sum_n \tilde{y}_{n,c,h_{m,c}}^2}{\sigma_z^2}+\frac{1}{s_\beta^2}\Big)^{-1}\Big\},
\end{aligned}
\tag{86}
$$

where the terms expressing the priors ($\pi(t_{m,c}|t_{m,-c})$, $\pi(g_{m,c}|t_{m,c})$ and $\pi(h_{m,c}|t_{m,c}, g_{m,c})$) are given by Equations 26, 28, 31, 32, and 33.

As already mentioned, a key point is to be able to sample directly from this conditional density in order to compute efficiently (i.e. avoiding repeated summing over $\mathcal{N}$) the terms corresponding to $t_{m,c}=1, g_{m,c}=1$ for all possible values of $h_{m,c}$. The computation of $\sum_n \tilde{y}_{n,c,h_{m,c}}^2$ does not necessitate summing over $\mathcal{N}$ since, from Equation 12, we have, in the case of a covariate of type "vector" where $h_{m,c}$ corresponds to the rank of the cut-off value used for binarisation,

$$
\sum_n \tilde{y}_{n,c,h_{m,c}}^2 \quad=\quad h_{m,c}\Big(\frac{h_{m,c}}{\mathcal{N}}-1\Big)^2+(\mathcal{N}-h_{m,c})\Big(\frac{h_{m,c}}{\mathcal{N}}\Big)^2 \tag{87}
$$

and, in the case of covariate of type "tree" where $h_{m,c}$ represents the index of a branch delineating a subtree whose size is denoted $|h_{m,c}|$,

$$
\sum_n \tilde{y}_{n,c,h_{m,c}}^2 \quad=\quad |h_{m,c}|\Big(\frac{|h_{m,c}|}{\mathcal{N}}-1\Big)^2+(\mathcal{N}-|h_{m,c}|)\Big(\frac{|h_{m,c}|}{\mathcal{N}}\Big)^2. \tag{88}
$$

Similarly, the sum $\sum_n \tilde{y}_{m,n,c,h_{m,c}}z_{m,n,-c}$ can be divided into two parts by separating the $y_{n,c}$ that are mapped to $\frac{h_{m,c}}{\mathcal{N}}-1$ and those that are mapped to $\frac{h_{m,c}}{\mathcal{N}}$. In the case of a covariate $c$

of type tree, the binarisation relies on the cut-off value denoted $y_{[h_{m,c}],c}$. Hence,

$$\sum_n \tilde{y}_{m,n,c,h_{m,c}} z_{m,n,-c} = \left(\frac{h_{m,c}}{\mathcal{N}} - 1\right) \sum_{n: y_{n,c} \le y_{[h_{m,c}],c}} z_{m,n,-c} + \left(\frac{h_{m,c}}{\mathcal{N}}\right) \left( \sum_n z_{m,n,-c} - \sum_{n: y_{n,c} \le y_{[h_{m,c}],c}} z_{m,n,-c} \right), \tag{89}$$

where the partial sums $\sum_{n: y_{n,c} \le y_{[h_{m,c}],c}} z_{m,n,-c}$ corresponding to all possible values of $h_{m,c}$ can be computed in one pass, by adding the $z_{m,n,-c}$'s in the order of increasing $y_{n,c}$'s. Similarly, in the case of a covariate of type "tree", all the relevant partial sums are computed in a single bottom-up recursion (i.e. taking the subtrees defining the binarisation in the order of increasing height as indexed in the rows of matrix $s_c$).

Importantly, the time complexity of the computation and sampling of the joint conditional of $t_{m,c}, g_{m,c}, h_{m,c}$ is therefore only $O(\mathcal{N})$, and not $O(\mathcal{N}^2)$ as a first look at Equation 85 might have suggested.
